# Why is human vision so poor in early development? The impact of initial sensitivity to low spatial frequencies on visual category learning

**DOI:** 10.1101/2022.06.22.497205

**Authors:** Omisa Jinsi, Margaret M. Henderson, Michael J. Tarr

**Affiliations:** Department of Psychology, Carnegie Mellon University, Pittsburgh, Pennsylvania, United States of America; Neuroscience Institute and Machine Learning Department, Carnegie Mellon University, Pittsburgh, Pennsylvania, United States of America; Department of Psychology and Neuroscience Institute, Carnegie Mellon University, Pittsburgh, Pennsylvania, United States of America

## Abstract

Humans are born with very low contrast sensitivity, meaning that developing infants experience the world “in a blur”. Is this solely a byproduct of maturational processes or is there some *functional* advantage for beginning life with poor vision? We explore whether reduced visual acuity as a consequence of low contrast sensitivity facilitates the acquisition of basic-level visual categories and, if so, whether this advantage also enhances subordinate-level category learning as visual acuity improves. Using convolutional neural networks (CNNs) and the ecoset dataset to simulate basic-level category learning, we manipulated model training curricula along three dimensions: presence of blurred inputs early in training, rate of blur removal over time, and grayscale versus color inputs. We found that a training regimen where blur starts high and is gradually reduced over time – as in human development – improves basic-level categorization performance relative to a regimen in which non-blurred inputs are used throughout. However, this pattern was observed only when grayscale images were used (analogous to the low sensitivity to color infants experience during early development). Importantly, the observed improvements in basic-level performance generalized to subordinate-level categorization as well: when models were fine-tuned on a dataset including subordinate-level categories (ImageNet), we found that models initially trained with blurred inputs showed a greater performance benefit than models trained solely on non-blurred inputs. Consistent with several other recent studies, we conclude that poor visual acuity in human newborns confers multiple advantages, including, as demonstrated here, more rapid and accurate acquisition of visual object categories at multiple hierarchical levels.

**Author Summary:** Why do humans start life with extremely poor vision? The common evolutionary story is that head size is small to accommodate the development of human-level intelligence. However, there is growing evidence that beginning life in a premature state confers short-term advantages. The “starting small” principle states that learning can be facilitated by restricted or impoverished inputs that reduce the learning complexity. We suggest that blurred vision during early development biases learning toward shape features, which organize objects naturally into “basic-level” categories that are the foundation of human cognition (e.g., birds, cars, etc.). Such categories are defined by their visual similarity in global object shape. Since blurring restricts visual inputs to emphasize shape information, it should enhance the learning of basic-level categories. We trained artificial neural-network models on basic-level categorization using either blurred or non-blurred color or grayscale images and found that models trained with blurred images outperformed models trained with non-blurred images, but only for grayscale. These same models performed better in subsequent “subordinate-level” tasks that required discriminating between objects within a basic-level category. Our simulations provide evidence that initially poor vision in infants has an important functional role in organizing knowledge about complex environments.

## Introduction

Why do human infants start life with such poor vision? Our altricial state at birth is often attributed to our extreme intelligence and the push “to be born even earlier to accommodate their [human infants] larger brains” (Piantadosi & Kidd, 2016). However, we posit that beyond providing an opportunity for avaricious corporations to sell “enrichment” toys to overachieving parents, low vision at the earliest stages of development facilitates the infant’s acquisition of basic-level visual categories. As an early learning objective, basic-level categories are at the core of the acquisition of stable mental concepts and foundational for naturally organizing large numbers of similar objects into behaviorally-relevant semantic clusters (e.g., “apple”, “table”, “fish”, etc.; Rosch et al., 1976). In their seminal paper, Rosch et al. (1976) propose that “Basic objects are the categories at the level of abstraction for which the cue validity of categories is maximized. Categories at higher levels of abstraction have lower cue validity than the basic because they have fewer attributes in common; categories subordinate to the basic have lower cue validity than the basic because they share most attributes with contrasting subordinate categories.” The question then becomes, what are the optimal set of visual attributes to form stable and functional categories? Rosch et al. (1976) suggest that the “shapes of objects” should exhibit a correlational structure reflecting high within-category similarity. Operationally, Rosch et al. define shape as the *outlines* or silhouettes of objects once their orientations have been aligned and their size normalized (Fig. 1A).

**Fig 1.**
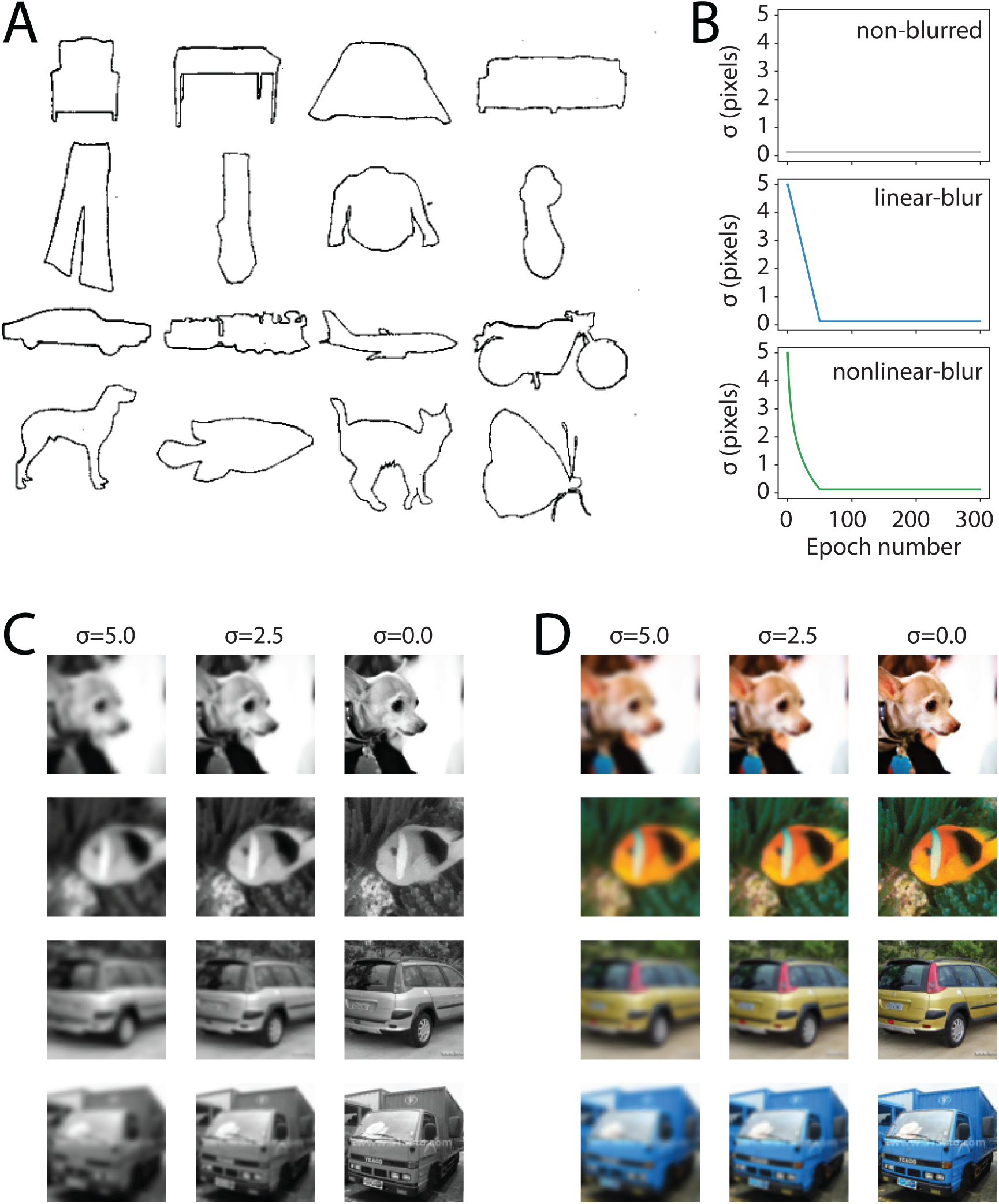
Diagrams depicting our experiment and rationale. **(A)** Rosch et al.’s stimulus images depicting exemplars from four basic-level categories within four superordinate categories (rows; adapted from Fig. 1, Rosch et al., 1976). **(B)** Schematics illustrating how the blurring parameter sigma (σ), in units of pixels, was varied over time in the three conditions of Experiment 1. In the non-blurred condition (top), sigma was fixed at a small value that results in no blurring throughout training, while in the two blur conditions, sigma started at 5 pixels and decreased over the first 50 epochs of training, corresponding to an increase in spatial acuity over time. See *Methods* for details. **(C-D)** Example ecoset images from the basic level categories “dog”, “fish”, “car”, “truck”, blurred at three example values for sigma (σ). Note that larger sigma values give rise to blurrier images, while σ = 0 results in an intact, unblurred image. **(C)** Examples of images in the grayscale condition. **(D)** The same images in the color condition.

Several subsequent studies lend support to the proposal that information about basic-level categories is carried by the coarse outlines/silhouettes of objects – which approximate the outputs of low spatial-frequency filtered versions of images. Cutzu and Tarr (1997) used a simple computational model of silhouette similarity and found that “views of objects from the same basic-level category are more similar to each other than to views of objects from different categories” (see also Gdalyahu & Weinshall, 1996). Inspired by this finding and related results, they suggest that human infants may perform binary basic-level categorization tasks (e.g., cats vs. dogs) based primarily on the information carried by object silhouettes. This prediction was borne out in Quinn, Eimas and Tarr (2001) who found that 3 and 4-month-old infants were able to form categorical representations for cats versus dogs based on object silhouettes. Although narrow in scope due to the use of only two categories and a simple pairwise discrimination task, this result does lend credence to the hypothesis that global outline shape may be critical for basic-level categorization. Reinforcing this point, French et al. (2002) observed that using input images filtered to remove high spatial frequencies – thereby emphasizing coarse outline shape over finer image details – improved an autoencoder’s ability to discriminate between the same cat and dog images as used in Quinn et al.’s (2001) study.

One concern is that in both of these studies, only a pairwise discrimination was tested: consequently, it is possible that the visual features supporting the discrimination between these particular 18 cats and 18 dogs do not generalize to the more complex space of generic basic-level categories. A second concern is that both studies used images of single objects against white backgrounds. As such, the discrimination task did not require visual object segmentation or figure-ground processing. That is, the stimulus images were inherently biased towards outline shape beyond what would be expected from natural images (which depict objects in the context of rich and complex visual environments; e.g., Figs. 1C-D). In sum, while these two studies are consistent with our argument, there is, to date, no robust demonstration as to whether information carried in the global shapes of objects is privileged with respect to acquiring basic-level category knowledge.

Stepping back, it is axiomatically true that the human visual system does not process images to extract the global shapes or silhouettes of objects. Our visual systems are built from a complex hierarchy of overlapping spatially-tuned neurons, the earliest of which respond to roughly circular regions of space and serve as *spatial frequency* filters (Hubel & Wiesel, 1959). Thus, as an approximation to global shape, a population of appropriately tuned neurons will produce a low-pass filtered image which will lack fine-grained details and which will highlight local regions of high contrast. Outline shapes or silhouettes similarly highlight global shape and contrast boundaries, but at the expense of *any* internal information. In this regard, using outline shapes or silhouettes as test stimuli further biases observers towards a singular aspect of object appearance in that object “interiors” lack even low-pass visual information.

In this context, we suggest that the basis of basic-level categorization within the human visual system is more plausibly anchored in images that are biased towards low-pass, high contrast information – a transform that emphasizes global shape and high contrast shape boundaries. Practically speaking, within our study we functionally approximate the visual experience of human infants by low-pass filtering input images – commonly referred to as “blurring”. As mentioned previously, human infant vision is blurry at birth. That is, it is much less sensitive to high spatial frequencies, but grows progressively more sensitive to higher spatial frequencies (and more adult-like) over the course of development (Dobson & Teller, 1978; Brown & Lindsey, 2009). More generally, as reviewed by Brown and Lindsey (2009), sensitivity to light, color, and contrast are all much lower in infants than as measured in adults. Underlying these limitations, infant contrast sensitivity is incredibly poor, measuring 50 times lower than adults at three months of age. Relevant to the manipulations used in our study, there is also evidence that the infant contrast sensitivity function is not only lower, but is shifted to lower spatial frequencies. Thus, at birth human infants experience a relatively blurry and colorless world that only improves slowly over the course of development – adult levels are not attained until at least three years of age.

From a theoretical perspective, we propose that poor vision at birth is not epiphenomenal. Rather, it is a functional constraint that provides a “leg up” for learning foundational knowledge about the world in the form of basic-level visual categories. Our proposal is an instance of the “Starting Small” principle put forward by Elman (1993). That is, initial information restriction in inputs may facilitate learning in terms of both rate of acquisition and ultimate performance. Intuitively, our argument is as follows. Basic-level categories form the conceptual “scaffolding” for much of our semantic knowledge (Rosch et al., 1976). Consequently, we assume that a core objective of early human development is to acquire robust basic-level categories – primarily through visual experience. Yet contrary to this goal, the visual world presents a complex, highly-detailed environment. While some attributes of this environment help to specify the basic-level structure of the world – through shared features across within-category exemplars – other attributes *detract* from learning this structure. In particular, fine-grained details of objects such as subtle shape variations and, especially for non-living things, colors or surface textures, often vary across category instances. As such, fine-grained features frequently *increase* the dissimilarity between items within a basic-level category. Thus, object information carried by high spatial frequencies may be detrimental to learning robust categories.

How then, does the infant learner select visual attributes appropriate to the learning objective? One possibility is that selective attention/inattention serves to orient the infant to low spatial frequencies in visual inputs. While such biasing might be theoretically achievable, it would require a great deal of “neural machinery” – a complex system devoted to identifying, orienting, and selecting across challenging visual inputs. In contrast (sic), the same end goal may be achieved in a straightforward manner by limiting contrast sensitivity at birth, but allowing it to improve over the course of development. Under this view, the limitation does not lie in the infant’s visual abilities *per se*, but rather in the way visual percepts are processed by the infant’s maturing visual system. Supporting this conjecture, Brown and Lindsey (2009) present evidence for a “critical immaturity that limits infant contrast sensitivity”. Moreover, they conclude that this limitation is a mid-level phenomenon and that “there is little effect of inattentiveness” in alert infants. As such, human infants are fully capable of visually exploring their surrounding environment (and, as a consequence, acquiring category knowledge), but their inputs are biased towards lower spatial frequencies, high contrast, and poor color perception due to intrinsic properties of their developing visual systems.

As antecedents to our present conjecture regarding basic-level category acquisition, the developmental trajectory of early vision has been shown to impact a variety of other visual domains. Notably, face recognition abilities have been found to interact with acuity in multiple ways. First, continually poor or unusually poor vision in infancy may also hinder the acquisition of adult-typical face recognition abilities (Maurer, Mondloch, & Lewis, 2007). For example, individuals who were treated for bilateral congenital cataracts early in life exhibit abnormal neural responses and deficits in face processing despite restored normal vision (Geldart et al., 2002; Putzar, Hötting, & Röder, 2010; de Heering & Maurer, 2014). Thus, a normal trajectory of improving contrast sensitivity (and associated other visual abilities) in early development is critical for normal visual recognition in adulthood, particularly for face individuation (Lewis & Maurer, 2009). Building on this finding, Vogelsang et al. (2018) tested the impact of blurred inputs during initial learning in a convolutional neural network performing face recognition. Similar to our hypothesis, they concluded that the “initial period of low retinal acuity characteristic of normal visual development induces extended spatial processing in the cortex that is important for configural face judgments”. That is, visual inputs that are blurry because of low acuity at birth facilitate learning critical spatial aspects of face processing that are essential for face identification abilities in adults. Supporting this conclusion, Jang and Tong (2021) likewise trained a convolutional neural network with images of both faces and objects that were initially blurred, but that, over training, became progressively less blurry. They found that the trained network’s recognition of faces, but not objects, was invariant over blur. Consistent with Vogelsang et al., Jang and Tong concluded that the holistic processing of faces benefits from initially poor vision in human infants.

Early experience with blurry images may also improve the robustness of visual recognition across image degradation. This is supported by a recent study by Avberšek et al. (2021), who used a coarse-to-fine image training regimen with multiple CNN models. More specifically, they attempted to mirror the trajectory of improving contrast sensitivity over early human development by initially training their models only with lower spatial frequency filtered images and then gradually introducing higher spatial frequencies as training progressed. As with other forms of invariance in CNNs, progressive training for a given perceptual dimension where variation is explicit (i.e., by isolating low spatial frequencies at the beginning of training) confers stronger invariance over that dimension. That is, Avberšek et al. found that models trained using a coarse-to-fine regimen performed significantly better on blurred images during validation testing. However, in contrast to the results we will discuss below, these benefits only maintained if blurred images were included throughout training. As such, invariance to image resolution may be learned differently in CNNs and in human infants.

Finally, as observed by Rosch et al. (1976), conceptual knowledge is hierarchical and includes finer-grained, or “subordinate” level, visual categories. As such, the human infant must ultimately acquire more than basic-level recognition skills. In particular, throughout the course of development and into adulthood, we also learn to differentiate between instances within basic-level categories. For example, imagine images of a Border Collie and a Siberian Husky. The general shapes at lower spatial frequencies of the two breeds of dogs are similar and, consequently, focusing on global shape information will facilitate their categorization as members of the same basic-level category, “dog”. On the other hand, differentiating between the two dog breeds necessitates drawing on details as carried by high spatial frequencies, for example fur texture, coloring, or more subtle shape differences such as snout size. Consistent with the conclusions above, improvement in contrast sensitivity and associated visual abilities, such as acuity, is essential for attaining adult-like category knowledge. Thus, the overall trajectory of visual development appears to be as functionally important as is the initial starting point.

Until recently it would have been impossible to address questions regarding different human developmental trajectories in any practical or ecological sense. While studies relying on recovered sight in older children or adults are somewhat informative, they are necessarily limited in their conclusions because of concurrent maturational changes that occur regardless of the structure of perceptual inputs (Maurer et al., 2007). However, the tools available for studying learning from experience have transformed over the last decade due to the rapid advance of artificial intelligence and computer vision in the form of deep convolutional neural networks (CNNs). Critically, CNNs have enabled the large-scale study of visual (and other domains of) learning at levels approaching human performance for many vision tasks (Bengio, LeCun, & Hinton, 2015). Thus, as a starting point, CNNs provide models that are high-performing approximations of human behavior for some tasks, for instance, object classification – whether they do so using the same computational principles as humans is an open question. One piece of evidence in favor of shared principles across artificial models and biological systems is that goal-driven CNNs trained on an object categorization task common to human visual behavior appear to learn object representations quite similar to *neural* representations of the same objects (Yamins & DiCarlo, 2016). That is, a wide array of studies have found that CNNs are able to account for much of the neural response variance in object viewing tasks as measured by fMRI in humans or by neurophysiological recordings in monkeys (Yamins et al., 2014; Kubilus et al., 2016). Given these similarities, CNN models provide an experimental setting in which to explore, using high-performing models and complex, real-world images, how category learning is affected by the manipulation of visual attributes such as blur or color.

In sum, we posit that poor infant vision at birth is not altricial by accident or for purely physiological reasons. Rather, consistent with past work, poor vision early in development may be a functional adaptation that bootstraps faster and more effective learning across multiple ecologically critical dimensions. We propose to examine this conjecture using a convolutional neural network trained to perform basic-level object categorization, using the ecoset database (Mehrer et al., 2020). Importantly, while prior studies exploring potential benefits have focused on face recognition, here we focus on general object recognition. Furthermore, our use of the ecoset dataset, as opposed to the popular ImageNet dataset which includes both basic- and subordinate-level categories, addresses a critical, currently unanswered question of how basic-level acquisition is impacted by early experience with blurred inputs. These experiments provide an assessment of how blurred images at initial learning impact the acquisition of basic-level visual categories – an organizing principle of conceptual knowledge that is central to adult cognition.

## Results

### Experiment 1

Models were trained to perform basic-level object categorization across 6 conditions defined by manipulations of spatial blur and color applied to training images drawn from the ecoset dataset (Mehrer et al., 2021; Figs. 1B-D). The spatial blur manipulation was intended to assess how the temporal dynamics of acuity changes across learning impact performance, by using two different time courses of blur reduction across training: blur decreasing linearly over time (linear-blur condition), and blur decreasing logarithmically over training (nonlinear-blur condition), in addition to a non-blurred image condition. The color/grayscale manipulation was motivated by the fact that color provides additional information with respect to object identity (i.e., some categories have highly consistent colors; Naor-Raz et al., 2003) as well as the known tendency of CNNs to be biased towards color and texture rather than shape (Geirhos et al., 2019). Both of these factors predict that CNNs trained with color images will have higher overall accuracy than those trained with grayscale images. In contrast, human vision is more shape biased and, critical to our hypothesis, human infants have poor color sensitivity (Brown & Lindsey, 2009). As such, our expectation was that we would observe stronger benefits for initial blurring in the absence of color.

As shown in Figure 2, both of these predictions were borne out. Across all spatial blur conditions, models trained on color images showed higher overall accuracy as compared to models trained on grayscale images (Fig. 2). Furthermore, the effect of blur on model performance differed between color and grayscale images. As shown in the left panel of Figure 2A, when using grayscale images, both the linear-blur and nonlinear-blur models achieved higher accuracy than models trained on non-blurred images, with linear-blur models performing slightly better than nonlinear-blur models. This difference between the linear-blur and non-blurred models manifested early in training, around epoch 25, and remained consistent throughout the remainder of training, while the difference between the nonlinear-blur and non-blurred models appeared later in training, closer to epoch 50, and was less consistent over subsequent training. No significant differences between the linear-blur and nonlinear-blur models were observed (linear mixed effects model with fixed effects of condition and epoch number, evaluated using a sliding window; significant effect of condition, FDR corrected ɑ=0.05; for details see *Methods).* Conversely, as shown in the right panel of Figure 2A, when using color images, there was little benefit for blurring training images, with all models converging to roughly the same level of performance by epoch 50.

**Fig 2.**
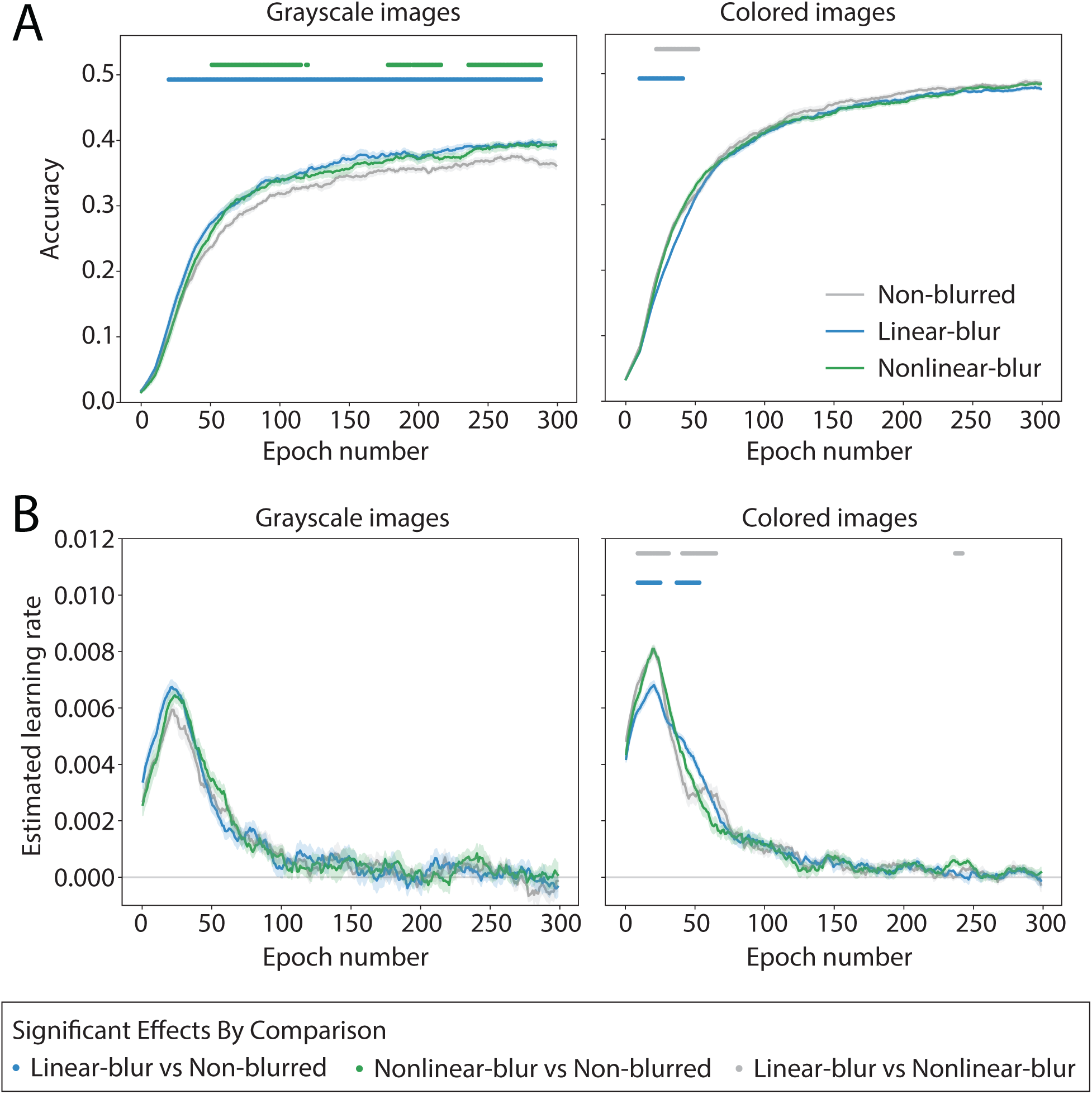
Experiment 1 model performance. Performance is shown over training for models in 6 conditions defined by the factors of blur and color. All models used the ResNet-50 architecture and were trained to perform basic-level object categorization using the ecoset image dataset. Each plotted point is an average across 10 runs of an otherwise identical model with different random seeds. Shaded error bars reflect mean ± SEM across these 10 trials. Dots along the top of each plot indicate time points at which a linear mixed effects model over a sliding temporal window revealed a significant effect of the specified pairwise condition comparison, in either direction (FDR corrected, ɑ=0.05). **(A)** Validation set accuracy averages. Gray corresponds to models trained with *non-blurred* images, blue to models trained with images whose blur decreases linearly over the first 50 epochs (*linear-blur* condition), and green to models trained with images whose blur decreases according to a logarithmic function over the first 50 epochs (*nonlinear-blur* condition). Left and right plots show averages for models trained using either grayscale images or color images, respectively. Accuracy was temporally smoothed to reduce noise. **(B)** Estimated learning rate computed as slope of accuracy over time. Colors correspond to models as in **(A)**.

To provide a more detailed picture of how the different conditions diverged over time, we computed estimated learning rates of each model as the approximate slope of accuracy over time. Learning rates peaked around epoch 25, then decreased rapidly, reaching near zero by the end of training. For the grayscale models, the linear-blur and nonlinear-blur models each achieved a slightly higher peak learning rate than the non-blurred model (Fig. 2B, left). Following this peak, the learning rate of the linear model fell slightly below the learning rate of the other two models, whereas the learning rate of the nonlinear-blur model remained marginally higher than both models. This short-lived dynamic matches the time at which the nonlinear-blur model was able to “catch up” with the linear-blur model in terms of validation accuracy. Finally, the learning rates of all grayscale models roughly converged by epoch 100. In the color models, the learning rate for the nonlinear-blur and non-blurred models both achieved a higher peak learning rate than the linear-blur models, but after this peak their learning rates decreased below the learning rate of the linear-blur model (Fig. 2B, right). The learning rates of all color models roughly converged by epoch 75.

### Experiment 2

The results of Experiment 1 indicate that, for grayscale images only, initial training with blurred visual inputs facilitates learning basic-level visual categories. In Experiment 2, we explored whether the improvements in learning categories at the basic level transfer to the acquisition of categories at the subordinate level. To address this question, we took the models trained in Experiment 1 and fine tuned them using a new image dataset, ImageNet, that included both basic-level and subordinate-level labeled categories.

As in Experiment 1, models pre-trained and fine-tuned with color images reached a higher overall accuracy level as compared to models pre-trained and fine-tuned with grayscale images (Fig. 3). In terms of the central manipulation in Experiment 2, for both color and grayscale images, there was a general benefit of pre-training on ecoset, with all pre-trained models, irrespective of the blur condition, showing higher validation accuracy at all time points as compared to models with no pre-training. This result is not surprising given that models trained from scratch would not have the benefit of previously learned features supporting basic-level categorization. More interesting are the observed differences in fine-tuning for grayscale models pre-trained with non-blurred images as compared to models pre-trained with blurred images (Fig. 3A, left). In contrast, few differences in fine-tuning were observed for color models in any pre-training condition (Fig. 3A, right).

**Fig 3.**
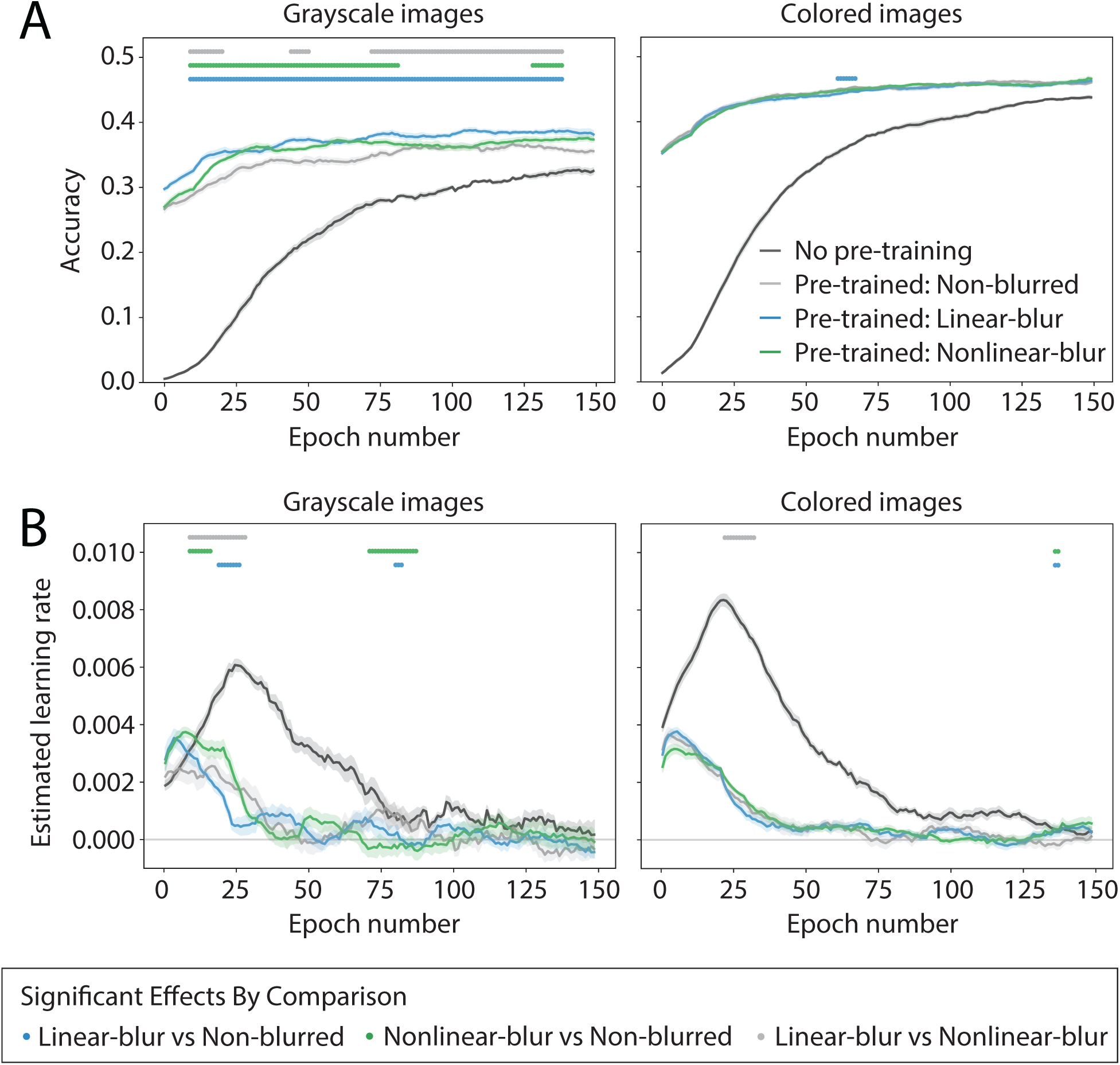
Experiment 2 model performance. Performance is shown for models from Experiment 1 that were fine-tuned with ImageNet images for 1000-way object categorization. ImageNet contains labels at different category levels, with some images labeled at the basic-level and others labeled at the subordinate-level. Fine-tuning was run only with non-blurred ImageNet images. Each plotted point is an average across 10 runs of an otherwise identical model with different random seeds. Shaded error bars reflect mean ± SEM across these 10 trials. Dots along the top of each plot indicate time points at which a linear mixed effects model over a sliding temporal window revealed a significant effect of the specified pairwise condition comparison, in either direction (FDR corrected, ɑ=0.05). **(A)** ImageNet validation set accuracy averages. Light gray corresponds to models trained with ecoset non-blurred images, blue to models trained with ecoset linear-blur images, green to models trained with ecoset nonlinear-blur images, and dark gray corresponds to new models with no pre-training, that is, starting from scratch. Left and right plots show averages for models trained using either grayscale images or color images, respectively. Accuracy was temporally smoothed to reduce noise. **(B)** Estimated learning rate computed as slope of accuracy over time. Colors correspond to models as in **(A)**. Only pairwise comparisons between different pre-trained models are shown; all pre-trained models performed significantly better than the no-pre-training models at all timepoints.

Because differences in pre-training conditions were only observed for grayscale models, the following discussion focuses only on results from the grayscale pre-trained models. Across many of the time points during fine-tuning, both the linear-blur and the nonlinear-blur models reached a higher validation set accuracy as compared to the non-blurred models. This difference was more pronounced for the linear-blur models, whose performance was significantly higher than the non-blurred model in all time windows. In contrast, the nonlinear-blur models were more variable, initially having a similar average validation accuracy as compared to the non-blurred models, but then increasing and approaching the accuracy of the linear-blur models after 30 epochs, and then briefly decreasing to the same accuracy as the non-blurred models. Overall, the accuracy of the nonlinear-blur models was significantly higher than the non-blurred models for roughly the first half of training, and again for several time windows near the end of the training interval. Finally, there was a significant difference between the linear-blur and nonlinear-blur models across several time windows, with better validation accuracy for the linear-blur models.

Pre-training with blurred images in a basic-level categorization task using ecoset benefits transfer performance for ImageNet, a dataset composed of both basic- and subordinate-level labeled images. One possibility is that this benefit is driven solely by the basic-level labeled images within ImageNet. Alternatively, representations learned during the acquisition of basic-level categories with blurred inputs may also confer some benefit for subordinate-level categorization. To address these possibilities, we hand-labeled the 1000 categories in ImageNet as either basic or subordinate and then re-computed the validation set accuracy for each of our grayscale models split by basic and subordinate labeled images. Figure 4 shows these results for each hierarchical category level as well as across all categories. Interestingly, differences in accuracy across pre-training conditions were similar for basic- and subordinate-level categories (as well as their aggregate). Again, not surprisingly, the three pre-training conditions showed significantly higher accuracy than the trained from scratch models. Most relevant to our current question, for both basic- and subordinate-level labeled images, linear-blur models showed significantly higher accuracy than either the non-blurred models or the nonlinear-blur models. This pattern is consistent with our earlier observation that linear-blur training with grayscale images produced the most robust advantages in basic-level categorization (Fig. 2A, left). Pertinent to the overall aims of Experiment 2, these benefits generalize from basic-level categorization to subordinate-level learning and categorization. Several factors may contribute to this transfer, including the initial learning of more robust basic-level categories or implicitly learned shape-biased subordinate-level categories acquired during the basic-level task.

**Fig 4.**
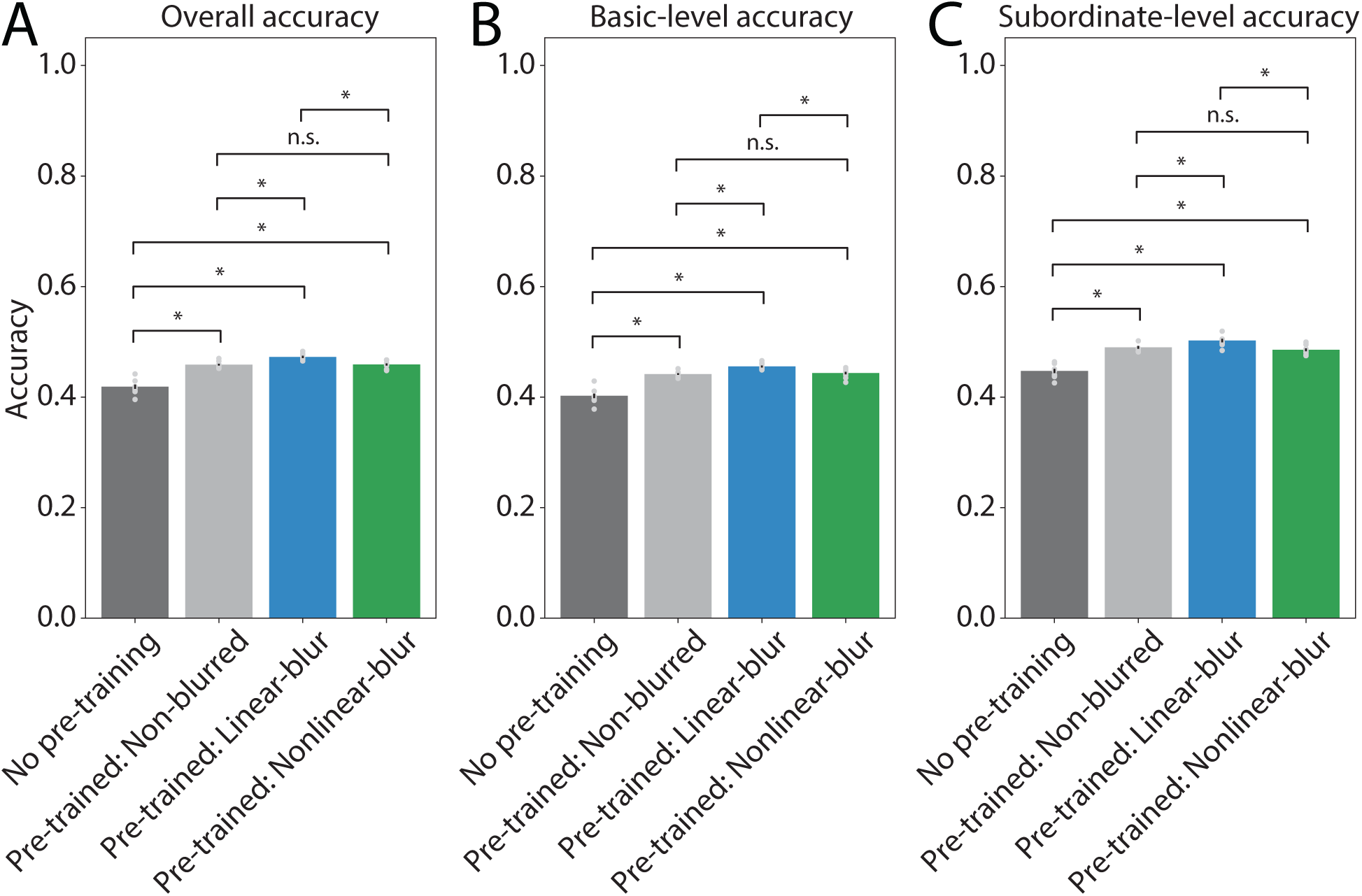
Experiment 2 model performance by category level. Performance is shown for ImageNet fine-tuned models pre-trained with grayscale images split by basic-level and subordinate-level category labeled images from ImageNet. Only results from models pre-trained with grayscale images are plotted because few differences were observed between models pre-trained with color images (Fig. 3). For grayscale pre-trained models, pre-training with blurred images confers similar benefits for basic-level and subordinate-level category labeled images. Validation accuracy was re-computed using all object categories and 1000-way categorization for **(A)** all categories, for **(B)** basic-level labeled object categories, and **(C)** subordinate-level labeled categories. Bar heights and error bars indicate mean ± SEM across 10 trials of each model, light gray dots show accuracy for individual trials. Brackets above bars indicate the significance of pairwise comparisons between conditions, assessed using a two-tailed independent samples *t*-test for each pairwise condition comparison (FDR corrected, ɑ=0.05) where * denotes a significant difference in either direction and n.s. denotes no significant difference.

## Discussion

Human newborns are strikingly helpless at birth – particularly relative to other primate and mammalian species. Piantadosi and Kidd (2016) speculate that a positive feedback loop has selected for intelligence whereby human infants’ premature state at birth enables the development of larger brains (and greater intelligence), but this same helplessness leads to a *need* for superior intelligence in adult caregivers. This account is compelling in terms of explaining long-term, evolutionary-scale benefits for altricial human newborns, but it fails to consider potential *near-term* developmental benefits that emerge as a consequence of human infant’s relatively underdeveloped cognitive and perceptual mechanisms. That is, additional selective pressures may have contributed to the initial state of human newborns and to the richness of adult-level conceptual knowledge.

Among a wide variety of desirable traits, superior human intelligence is characterized by complex semantics across thousands of categories (Huth et al., 2012). This rich representation of conceptual knowledge is grounded in basic-level categories (Rosch et al., 1976) and their rapid and stable acquisition in early development is central to building a robust semantic foundation. Under this view, we hypothesize that the underdeveloped state of the newborn human visual is functional and, among other things, facilitates the infant’s rapid acquisition of basic-level categories. The work we present here, along with several related studies (French et al., 2002; Vogelsang et al., 2018; Avberšek et al., 2021; Jang & Tong, 2021), supports this hypothesis, suggesting that poor infant vision – in the form of low contrast sensitivity – confers multiple adaptive benefits to the visual learner.

Within both our current computational experiments and many of these related studies, underdeveloped human newborn vision was simulated by restricting visual inputs to low spatial frequencies (“blurring”). This transform emphasizes global shape – particularly in an object’s bounding contours – because fine-grained shape details and textures are selectively removed from the image. As such, the resultant inputs emphasize the kinds of visual information on which basic-level categories are based (Rosch et al., 1976) and we predicted that the learning of such categories would be enhanced when training with blurred images relative to training with full-resolution images.

Our prediction was confirmed across both of our experiments. In Experiment 1, training on blurred images improved the accuracy of learning for basic-level categories relative to training with non-blurred images. In Experiment 2, pre-training on blurred images in a basic-level categorization task transferred to and improved the learning of subordinate-level categories relative to pre-training with non-blurred images. However, in both experiments, these benefits only manifested when using grayscale, and not color, images. In contrast to human perceivers, visual CNNs appear to be biased towards relying on color and texture more than shape (Geirhos et al., 2019; Hermann, Chen, & Kornblith, 2020). As such, shape transforms such as blurring may have had less effect on model performance in the presence of fine-grained shape and texture information. That is, when models have access to color and high spatial frequency information, they may learn on the basis of these kinds of features at the expense of other features such as global shape. However, one might reasonably ask why color and high spatial frequency information should be excluded from initial training given that such models show better overall performance?

There are several reasons for this. First, color is quite variable across many, if not most, object categories – only a handful show reliable, diagnostic colors (Naor-Raz et al., 2003) – so it may not be in a learner’s best interest to attend to color when it provides negative information for the majority of categories. Second, as just noted, humans and CNNs learn high-level visual representations with different feature biases. Thus, removing color may bring CNNs into better alignment with human biases, particularly with infants who are generally more dependent on global outline shapes than surface features (Landau et al., 1992). Third, low contrast sensitivity in human infants not only affects spatial resolution but also color sensitivity (Brown & Lindsey, 2009). As such, the learning environment for infants deemphasizes color and CNNs may better approximate the experience of human infant learners when color is absent. Moreover, these same biases in early development may help to account for the shape bias observed in human adults. Returning to the first of these reasons, the variability of color for many basic-level categories also points to another potential benefit of low contrast-sensitivity in infants – color may actually harm the acquisition of stable and generalizable basic-level categories. Finally, although beyond the scope of our current work, we note that the color and texture biases for CNNs as well the overall higher performance we observed for color image trained models may reflect representational capacity advantages for CNNs over humans. That is, while both humans and CNNs appear to learn generalizable models for visual categories, CNNs appear to be better able to include, with basic-level representations, specific instances of information that is only useful for a subset of the category.

Note that the benefits of training with blurred images only manifested under certain specific model training conditions. In particular, using a learning rate scheduler (which sets step size for weight updates) in Experiment 1 rendered the effect of blur less apparent as compared to when the scheduler was removed. We speculate that using the scheduler may have optimized learning to the point that the rapid shift to extremely high performance may have resulted in an environment in which there was little room for blurring to further improve performance. In contrast, human infant categorization performance is quite poor and only slowly improves – over many years – to adult-like levels. One interesting possibility that should be pursued in a new study is whether models using a learning rate scheduler, despite high performance, learn less robust basic-level category representations.

Another model parameter that had some impact on performance was an advantage for the linear-blur condition over the nonlinear-blur condition. In the former condition the degree of blur (i.e., standard deviation of Gaussian filter) decreased linearly over time, while in the latter condition the degree of blur decreased more quickly at the start of training. Given that the standard deviation of the filter in the spatial domain is nonlinearly related to its standard deviation in the frequency domain, the nonlinear-blur condition may more closely approximate a linear increase in acuity than does the linear-blur condition. Based on the finding that acuity increases approximately linearly across human visual development (Courage & Adams, 1990), we had initially hypothesized that our nonlinear-blur models would achieve higher performance than our linear-blur models. However, our results showed the opposite. There are multiple plausible explanations for such a result. First, CNNs may benefit more from a linear time course of blur reduction because neural networks follow somewhat different learning principles than humans. Second, it is also possible that our implementation of nonlinear blur was too fast relative to the time period in which we implemented blurring of images. In preliminary tests, we found that our slowest nonlinear function (with a base of 2) was the most effective out of the three tested nonlinear functions. Although it was the best out of the three, given the brief span in which the model was trained on blurred images, it is possible that the decrease in blur was still not sufficiently gradual for the network to benefit from the biasing provided by blurring. Future work should explore models with slower nonlinear blurring functions as well as training on blurred images for longer periods of time in order to evaluate how these parameters affect performance in nonlinear models.

Interestingly, the results of Experiment 2 suggest that the benefit of early experience with blurred images generalizes to subordinate-level categories. Specifically, models initially trained to perform basic-level categorization on ecoset showed a boost in ImageNet accuracy over randomly initialized models, and that this benefit was largest for models trained with blurred images. Critically, this benefit was observed for both basic-level labeled images *and* subordinate-level labeled images from ImageNet. Given the assumption that subordinate-level categories are defined more by fine-grained shape and surface appearance (Rosch et al., 1976), why does a model trained on blurred images facilitate learning subordinate-level categories? We observe that there is some similarity in the representations needed for basic-level versus subordinate-level classification in that subordinate-level categories are typically refinements of basic-level categories. To the extent that this is true (e.g., most dog breeds are all visually recognizable as dogs), the basic-level representations learned by the linear-blur and nonlinear-blur models in Experiment 1 can be efficiently fine-tuned for a new subordinate-level task in Experiment 2. However, in the case of basic-level categories with subordinate-level members that deviate from the general category appearance (e.g., penguins and ostriches do not look like the majority of birds), our expectation is that fine-tuning for such subordinate-level categories will not benefit from pre-training with blurred images. As future work, fine-grained analyses of category learning that consider the specific visual and semantic structures for individual categories will help us to better understand how early experience impacts category learning at multiple hierarchical levels.

Related to our observation that basic-level pre-training benefits later learning of subordinate-level categories, a recent study found that training a visual CNN model with superordinate-level labeled images (e.g., fruit, animal) followed by basic-level training resulted in a model that was highly robust to image perturbations, as well as exhibiting a stronger shape bias as compared to models trained without hierarchical labels (Ahn et al., 2021). In tandem with our results, these findings highlight the importance of considering the role of different hierarchical levels when creating training paradigms for neural networks. For example, ImageNet includes a mixture of basic-level and subordinate-level labels that are not meaningfully differentiated, thereby confounding two levels of category learning that are likely to be supported by different visual features. At the same time, some ImageNet categories labeled at the subordinate-level may effectively function as basic-level categories in that no other subordinate-level categories for their parent basic-level category are included in ImageNet. Within the work we presented here, we attempted to control for some of these issues in Experiment 1 by using the ecoset dataset (Mehrer et al. 2021). Critically, ecoset contains only basic-level labels and is motivated by the foundational role that the basic level plays in category learning and adult cognition (Rosch et al., 1976). Moreover, when using ImageNet in Experiment 2, we explicitly split our results by basic- and subordinate-level labeled images. Future work should also explore superordinate-level category labels to investigate how early experience with blurred inputs interacts with a hierarchical superordinate-to-basic training regimen similar to that used by Ahn and colleagues.

Finally, several recent studies have also investigated how training with blurred images impacts learning in neural network models. Avberšek et al. (2021) investigated the effects of training a neural network on various regimens of progressively blurry-to-clear images. They were particularly interested in how the training regimen impacts a neural network’s processing of high and low spatial frequencies, including robustness to blur. Models whose training incorporated both blurry and non-blurred images were better able to categorize low-pass filtered versions of images than were models whose training included only non-blurred images. At the same time, contrary to our results, Avberšek et al. reported, for categorization of un-filtered images, that models trained solely on non-blurred images always performed better than models whose training incorporated blurred images. Similarly, Jang and Tong (2021) compared the effect of blurry-to-clear training on face and object recognition did not observe an overall benefit in categorization for blurry-trained models over non-blurred models. While the models used in these studies and our present study varied across many model parameters, one plausible explanation for the discrepancy between their studies and ours is that we utilized the ecoset image database, which includes only basic-level categories, while both the Avberšek et al. and Jang and Tong studies used the ImageNet image database. As discussed above, the basic-level holds a privileged position in human cognition and may be best captured by features different from those best suited to subordinate-level tasks (Rosch et al., 1976). In particular, we suggest that blurred inputs only confer a benefit for tasks requiring basic-level categorization in that this image transform biases inputs specifically towards the kinds of information theorized to be critical to the structure of most basic-level categories. We also note that a particular set of model parameters were necessary for the effect of blur to be detected. These factors, along with other aspects that differed across studies, for example, network architecture, may have led to different outcomes. However, when experimental conditions are specifically targeted at mirroring the learning environment of the human infant – not only reducing image resolution, but also removing color, and restricting the task to basic-level category learning – we find a clear benefit for models whose training includes blurred inputs at initial learning.

In conclusion, our computational simulations support the theory that low visual acuity in early development (as a consequence of low contrast sensitivity) is a key factor in infant visual growth and cognitive development, providing an early advantage in basic-level category learning. Importantly, although blurry inputs were only presented briefly at the start of training, early performance advantages were sustained throughout the duration of basic-level training and persisted through the introduction of subordinate-level categorization tasks. Thus, poor vision early in life, rather than hindering learning, is a functional adaptation that supports the human infant’s acquisition of robust conceptual structures.

## Materials and Methods

Our study can be split into two distinct sections. First, Experiment 1, using standard CNNs, explores whether using blurred visual inputs during the initial learning of basic-level categories enhances either the rate of learning or the attained accuracy in categorization. Second, Experiment 2, using the same CNNs trained in Experiment 1, explores whether the basic-level representations learned with initially blurred visual inputs will also confer benefits to learning subordinate-level categories.

### Models and Datasets

For all models in both experiments, we used the ResNet-50 architecture (He et. al., 2016) with a learning rate of 0.1 and a standard gradient descent (SGD) optimizer. Models were trained using PyTorch version 1.10.0 in Python version 3.7.1, on the Carnegie Mellon Neuroscience Institute High Performance Computing Cluster which consists of 21 CPU nodes and 12 GPU nodes, 280TB terabytes of shared disk space and 2.8 terabytes of RAM (https://ni.cmu.edu/computing/knowledge-base/computing-facilities-description-overview/ ). For our SGD optimizer, we set momentum to 0.9 and weight decay to 0.1 . No learning rate scheduler was used, a decision motivated by our finding in initial tests that when a scheduler was used, all our models performed similarly regardless of the amount of blur applied to images; we discuss the implications of this in our *Discussion*. Individual models were each trained for 300 epochs (Exp. 1) or 150 epochs (Exp. 2). To train our models, we utilized the ecoset image dataset which contains over 1.5 million images drawn from 565 labeled basic-level categories (Mehrer et al., 2021) - this resulted in a final output layer of 565 units. In contrast, the more common ImageNet image dataset (Deng et al., 2009) contains a mixture of labeled basic- and subordinate-level categories (e.g., bird species and dog breeds). Precisely because of this mixture of labeled categories, ImageNet was used for fine-tuning our models in Experiment 2. To address images in ecoset and ImageNet being of variable sizes, we used the PIL image processing library in Python to resize each image by center cropping based on the minimum dimension between width and height and then resized all images to 224×224 pixels (code available at https://github.com/ojinsi/startingblurry).

### Experiment 1

To explore the impact of blurring during learning, we trained multiple models on image sets defined by different numbers of blurred and non-blurred training images. All models were trained on the same 50,000 randomly selected images per epoch, but with different amounts of blur applied to the training images during pre-processing. For validation, for each epoch, we computed validation set accuracy using the entire ecoset validation set of 28,245 images. Images in the validation set for a given epoch were blurred at the same level as were the training images for that epoch. The use of 50,000 images per epoch was selected based on a balance between having a sufficient number of images to adequately train the model and limiting the number of images to allow gradual blur to have some impact on learning. For all blurred models, images were blurred for only the first 50 epochs. Pilot testing using blurring for the first 50, 100, 150, 200, 250, or 300 epochs in different models revealed that limiting blur to the beginning of training – 50 epochs – produced our most robust results.

Blurring was realized by using a Gaussian filter, implemented using the GaussianBlur function in the transformations module under Torchvision (a library of Pytorch). This filter applies a low-pass filter to each image by removing any spatial frequencies finer than the scale of the Gaussian (Figs. 1C-D). Implementing a Gaussian blur filter requires calculating sigma (*σ*, which denotes the standard deviation of the Gaussian filter, in pixels) and a kernel size value based on the epoch. In all blurred models, initially *σ* = 5 and was reduced until the 50th training epoch, at which point *σ* = 0.25 . Our implementation and manipulation of Gaussian blur is similar to the manipulations used in past work on how image blur impacts different aspects of visual learning (Jang & Tong, 2021; Vogelsang et al., 2018). In different models we used either a linear or a logarithmic function to determine the how much the value of *σ* was reduced for each subsequent epoch. The linear function was defined by calculating the difference between our initial and final *σ* values and dividing by 50, the number of epochs in which images were blurred. The value of *σ* was then decreased by this calculated constant after each epoch. The logarithmic function was defined by reducing the value of *σ* according to a logarithmic function with a base of 2, which results in a reduction in *σ* which is larger at the beginning of training (Fig. 1B). Since the *σ* of our Gaussian kernel is related nonlinearly to the resulting low-pass frequency cutoff of the image (our approximation of visual acuity), a logarithmic change in *σ* means that the change in acuity over time will more closely approximate a linear function, as has been measured from human developmental data (Courage & Adams, 1990). For both functions, kernel size was calculated as 8 times the *σ* value plus 1 to ensure that: 1) the kernel size was an odd number; 2) the entire kernel was sufficiently large so as to accommodate ±4 standard deviations from the center of the filter.

In addition to the amount of blurring applied to each image, we also manipulated the color content of images by using either full-color or grayscale images. Our rationale for manipulating color was two-fold. First, poor contrast sensitivity in infants also limits their ability to see color differences (Brown & Lindsey, 2009). Thus, images absent color in addition to image blur may better approximate a human infant’s visual experience in the early months of their development. Second, while color can play a role in human categorization (Naor-Raz, Tarr, & Kersten, 2003; Kimura et al., 2010), color is not consistent or diagnostic for many basic-level categories. To the extent that CNNs tend to overfit, fine-grained details such as color or texture (e.g., Geirhos et. al., 2019) may support category learning at the expense of more general shape properties. We hypothesized that models trained with grayscale images might show a stronger shape bias and, consequently, lead to a larger impact of spatial frequency manipulations on learning.

In total, we ran six model conditions: three blur conditions, *non-blurred* (normal unfiltered inputs throughout training), *linear-blur* (a linear decrease in blur over the first 50 training epochs), and *nonlinear-blur* (a logarithmic decrease in blur over the first 50 training epochs), crossed with two color conditions, *color* and *grayscale* images. For each of these six conditions, we ran 10 replicates (“trials”) of the otherwise identical model with different random seeds. To ensure comparability of our results across conditions, we used the same random images for epoch during training and the complete validation set after each training epoch (although the appearance of each image in terms of blur and color differed based on the condition).

For the purpose of visualization and statistical analyses, we temporally smoothed the validation accuracy results for each individual model, by computing the moving average over a sliding window of 20 epochs. All subsequent statistical analyses were then performed on this smoothed data. Learning rate was then estimated based on this temporally-smoothed data by finding the difference in validation accuracy between neighboring epochs (i.e., the approximate slope of the accuracy), and then applying a second moving average filter with a window size of 20 epochs. For both the accuracy and learning rate values, we then tested for significant differences between conditions at each timepoint by using a linear mixed effects model, implemented using the Python package *statsmodels.* Specifically, we used a sliding window where 20 epochs were considered at a time, and we constructed a model where condition and epoch number were modeled as fixed effects (categorical and continuous, respectively), while the trial number was modeled as a random effect. We used this analysis to obtain a coefficient and *p*-value for the effect of condition, for each possible pairwise condition comparison (non-blurred vs. linear-blur, non-blurred vs. nonlinear-blur, linear-blur vs. nonlinear-blur). Comparisons were always done between different blur conditions within the same color condition; no direct statistical comparisons were done between the grayscale and color models. Finally, the resulting *p*-values from all pairwise comparisons, as shown in Figure 2, were FDR corrected across all epochs using the Benjamini-Hochberg procedure implemented in *statsmodels*, with ɑ=0.05 (Benjamini & Hochberg, 1995).

### Experiment 2

To explore whether the benefits conferred by using blurred training images for basic-level categorization generalize to subordinate-level recognition, we used a transfer learning paradigm in which the trained models from Experiment 1 were fine-tuned using ImageNet (Deng et. al., 2009). ImageNet was used in for Experiment 2 because, in contrast to ecoset, ImageNet is composed of both basic- and subordinate-level category labels. The overall design of Experiment 2 was intended to maintain consistency with Experiment 1. As in Experiment 1, 50,000 images were randomly selected for training per epoch and all models were trained using these same images. ImageNet images, which are of variable sizes in their raw form, were resized using the same center crop method as used in Experiment 1. Color content was held constant when generalizing from ecoset to ImageNet: when using an ecoset pre-trained model that was initially trained with grayscale images, all ImageNet fine-tuning was performed with grayscale images, and vice versa for color models. To maintain a similar learning environment as in Experiment 1, models in Experiment 2 used the same architecture and hyperparameters, except for two differences. First, because Experiment 2 is based on fine tuning (and not training from scratch), models were trained for only 150 epochs (rather than 300 epochs). Second, because of the larger number of classes labeled in ImageNet, models used for fine-tuning in Experiment 2 had a final layer with 1,000 units (rather than 565 units).

To determine which models would serve as the base for Experiment 2, we identified the best training time point (based on validation accuracy) for each model from Experiment 1, and the weights from the model at this time point were stored (note that this could be a different time point for different trials in a given condition). In Experiment 2, these stored models were loaded into our adjusted architecture and then fine-tuned with ImageNet images. In addition to these stored models, we also trained, from scratch with random initial weights, two control models with either grayscale or color images. This resulted in eight total models (six pre-trained models and two control models). As in Experiment 1, we ran 10 trials for each condition. For each trial in the pre-trained model conditions, the pre-trained model was from the corresponding trial in Experiment 1. For example, during trial 4 of Experiment 2, the starting point in each condition was trial 4 of the corresponding model from that condition in Experiment 1.

To calculate validation accuracy and learning rate, we used the same averaging techniques as in Experiment 1. Statistical tests comparing training conditions were also identical to the analysis procedure used in Experiment 1. After running all trials for grayscale models we also performed an analysis in which we computed the validation set accuracy separately for the ImageNet categories that were defined as “basic-level” versus those defined as “subordinate-level”. We focused on the models trained with grayscale images only because these models showed the largest effect of blur condition. ImageNet categories were defined as basic or subordinate categories based on a manual estimate of how frequently the category name would be used in everyday language to refer to the object of interest (Rosch et al., 1976). For example, “bee” and “strawberry” were labeled as basic-level, while “Yorkshire terrier” and “Granny-smith apple” were labeled as subordinate-level. The full list of basic- and subordinate-level assignments for the 1000 ImageNet categories can be accessed in our supplementary materials (Supplementary Tables 1-2).

To compute accuracy separately for the basic and subordinate categories, we first identified the best training time point with respect to validation accuracy for each individually fine-tuned model in Experiment 2, and saved the weights for this best time point for each model. We then ran the entire ImageNet validation dataset through the models with these saved weights, and computed accuracy for one label at a time. Accuracy was defined as the number of correctly classified images of a chosen label divided by the number of images of the chosen label. The resulting category-specific accuracy values were then averaged over either all basic-level categories or all subordinate-level categories. For comparison, we also computed the overall validation accuracy of each model across all categories. To facilitate comparison of this value with the basic- and subordinate-level accuracy values, in computing overall validation accuracy we always used the same time point as was used to generate the basic- and subordinate-level accuracy values (i.e., the time point for each model with the single best validation accuracy value). Finally, we performed statistical comparisons between the four different pre-training conditions (no pre-training, pre-training with non-blurred images, pre-training with linear-blur images, pre-training with nonlinear-blur images) using two-tailed independent samples *t*-tests between each pair of conditions, implemented using the Python package *scipy*. The resulting *p*-values from all pairwise comparisons, as shown in Figure 3, were FDR corrected as described above.

### Code Availability

All code needed to reproduce our experiments and analyses is publicly available on GitHub, at https://github.com/ojinsi/startingblurry.

## Acknowledgements

^†^The first two authors contributed equally to this work. MMH was funded by a Distinguished Postdoctoral Fellowship from the Carnegie Mellon Neuroscience Institute. The authors thank the following people for contributing ideas and commentary to this project: Jayanth Koushik.

## SUPPLEMENTARY MATERIALS

**Supplementary Table 1.**
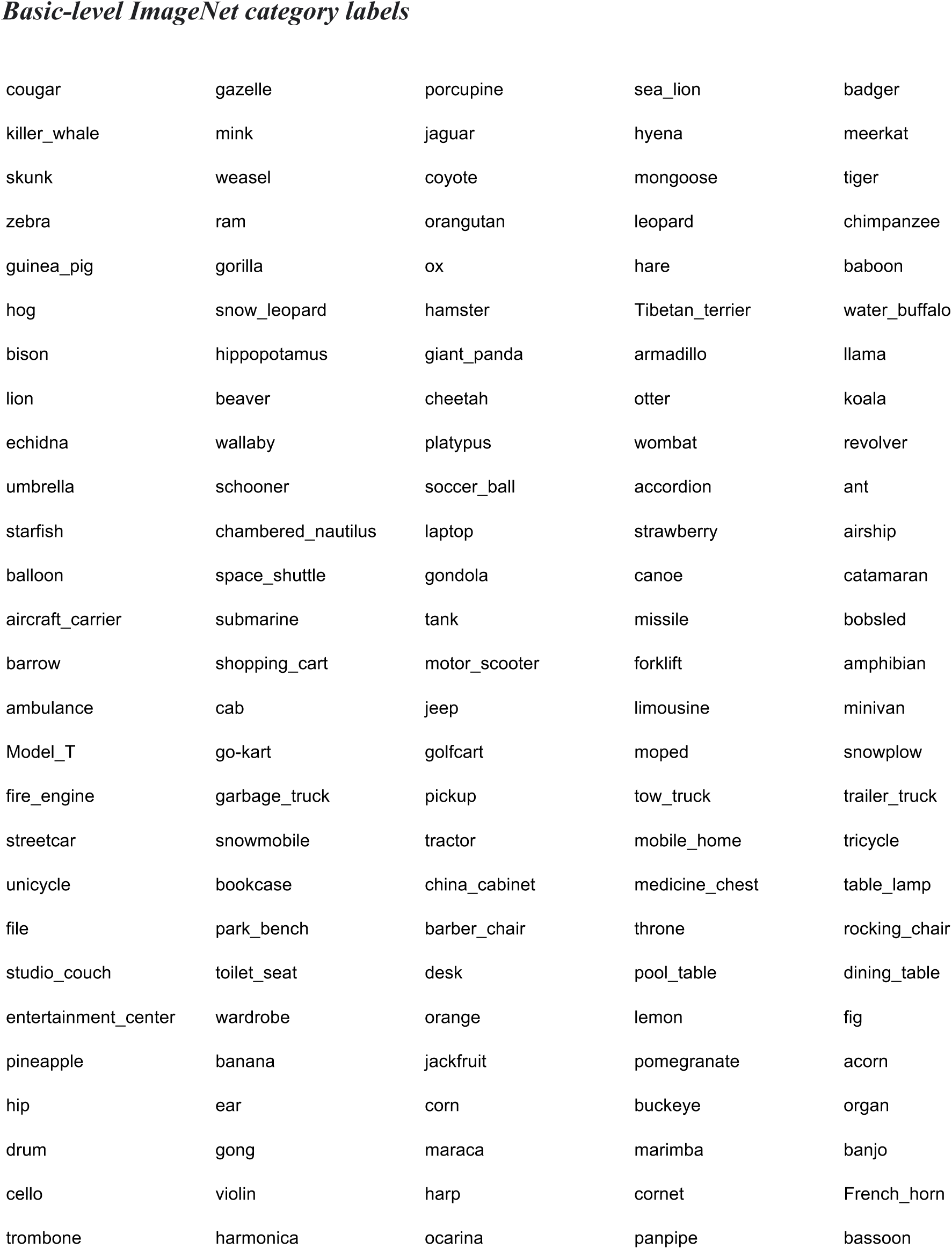
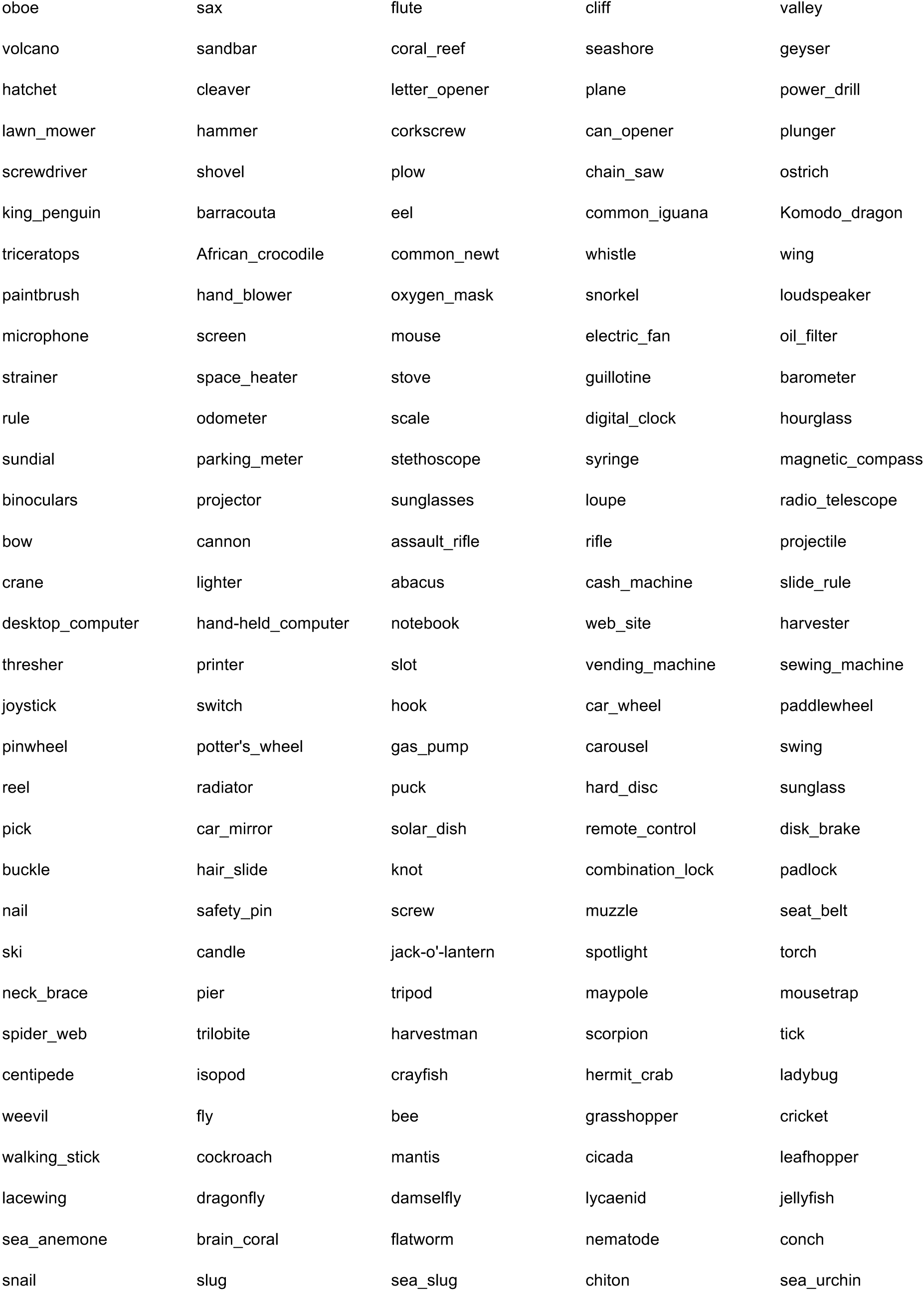
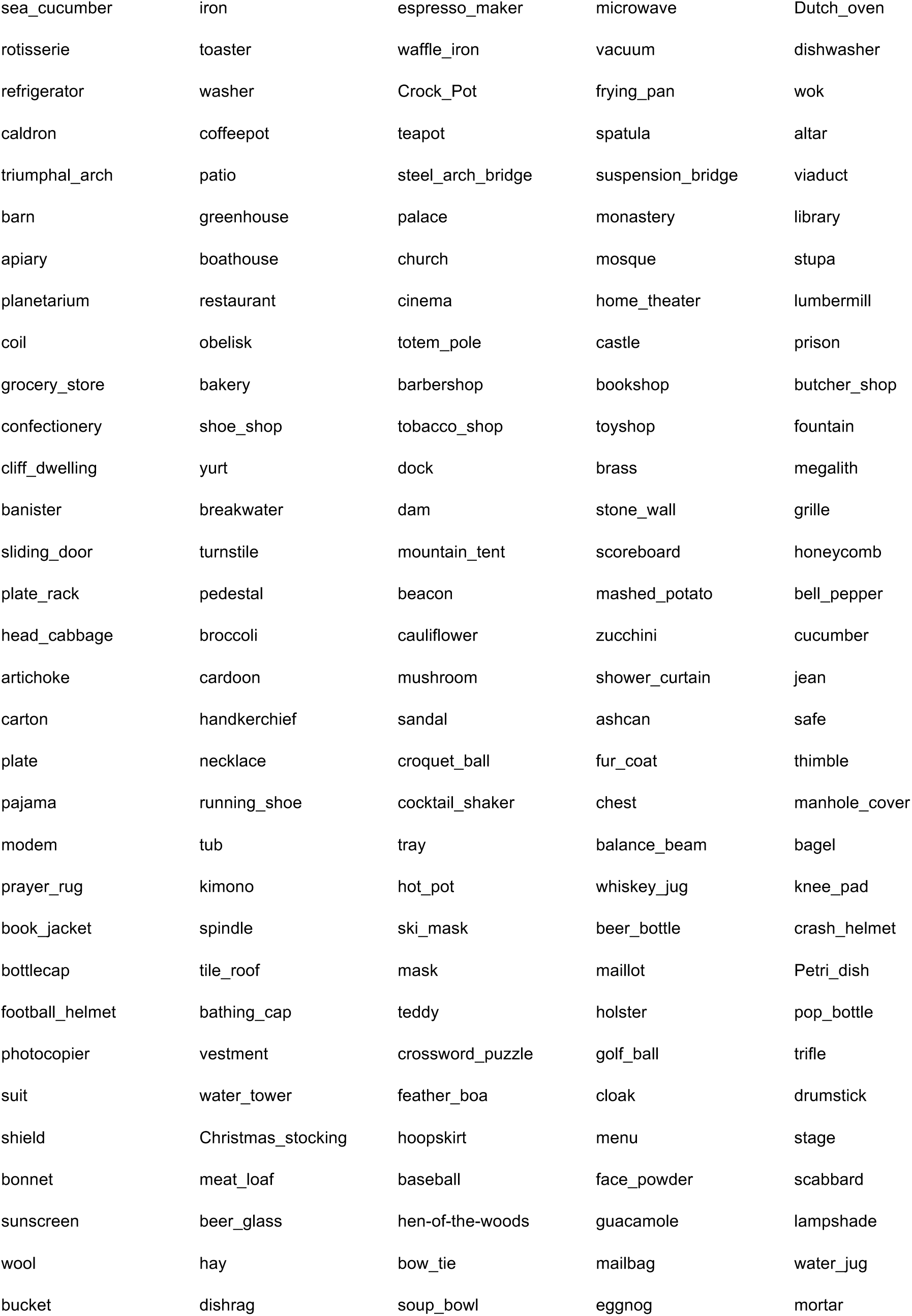
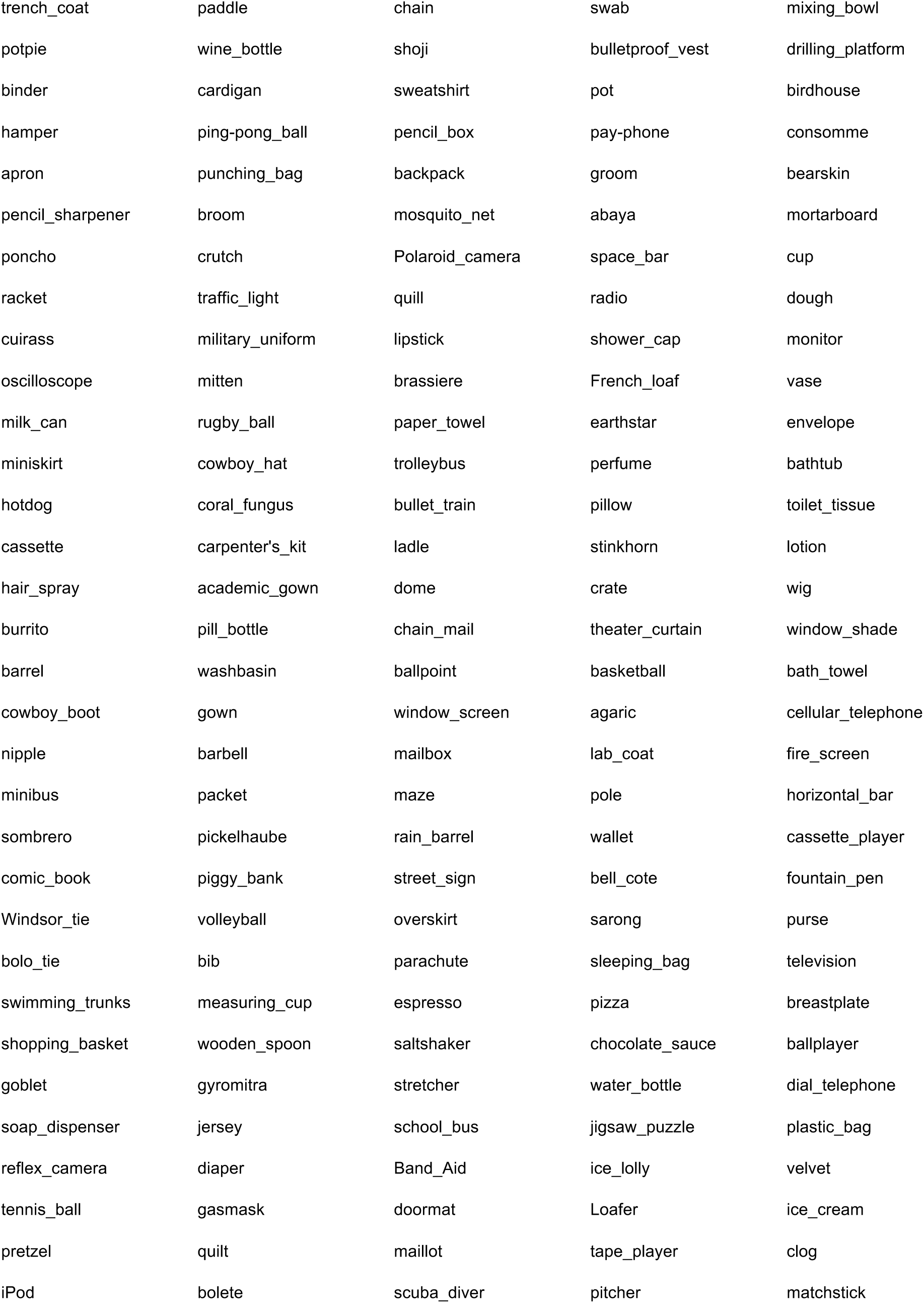
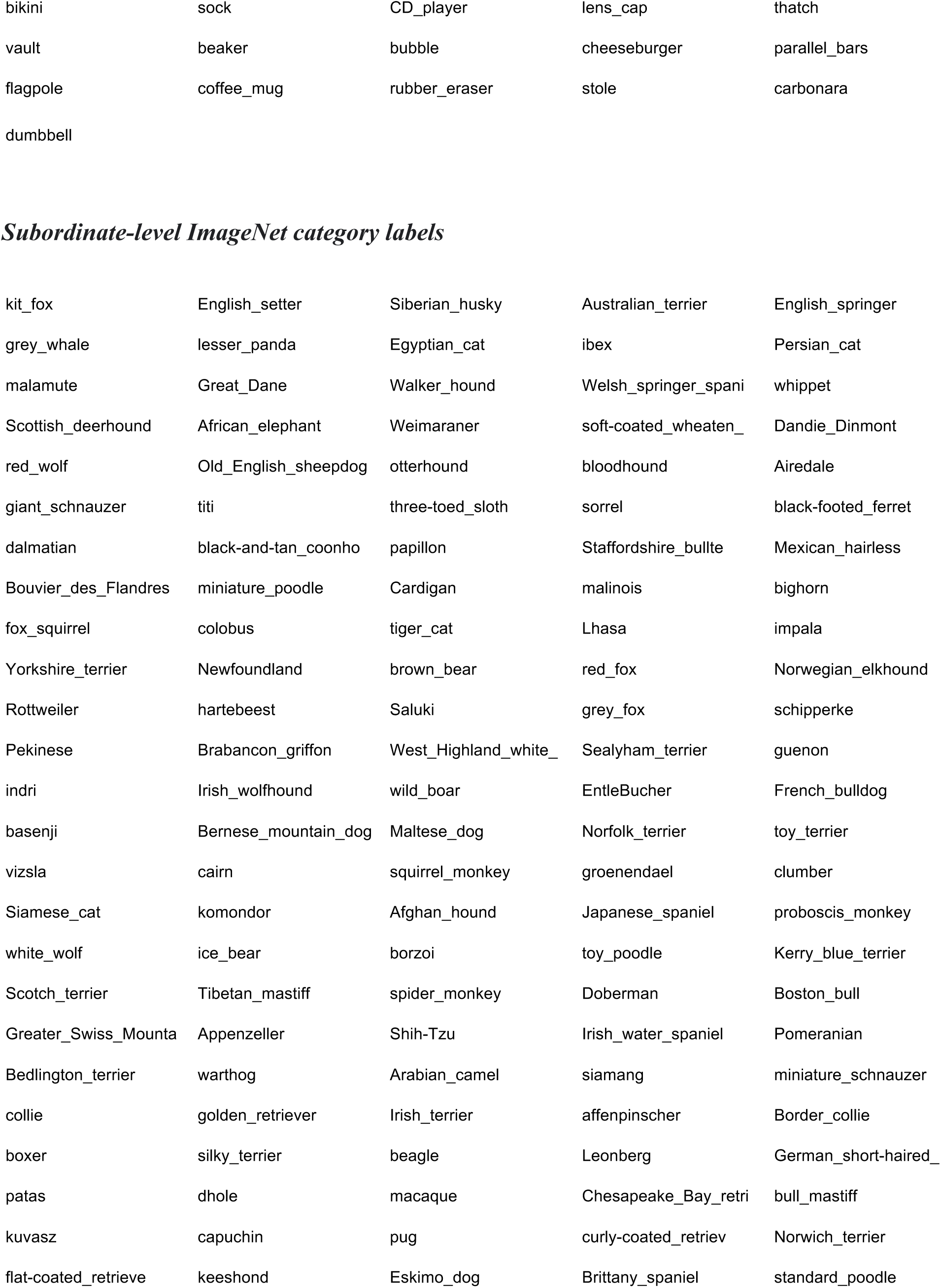
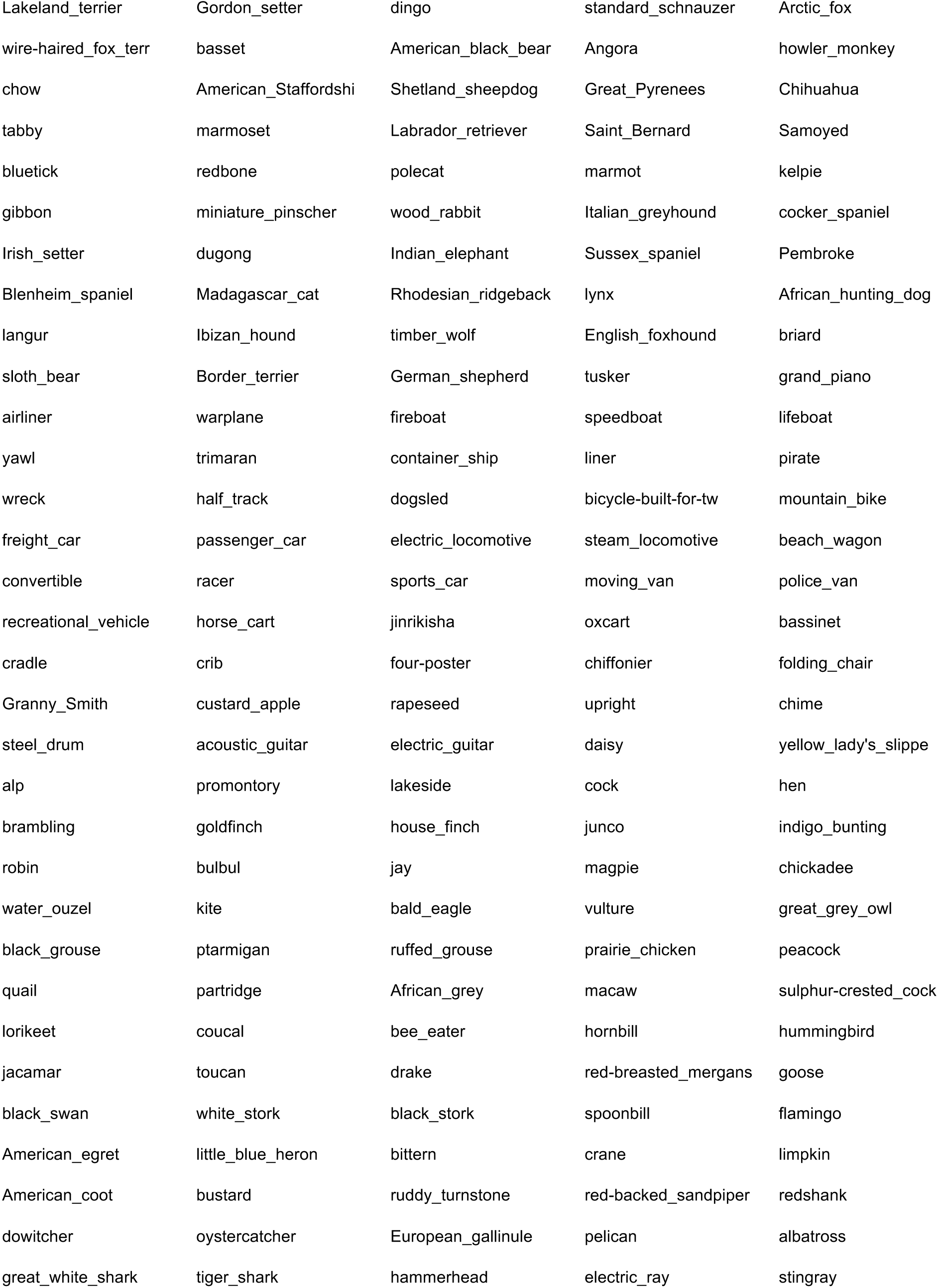
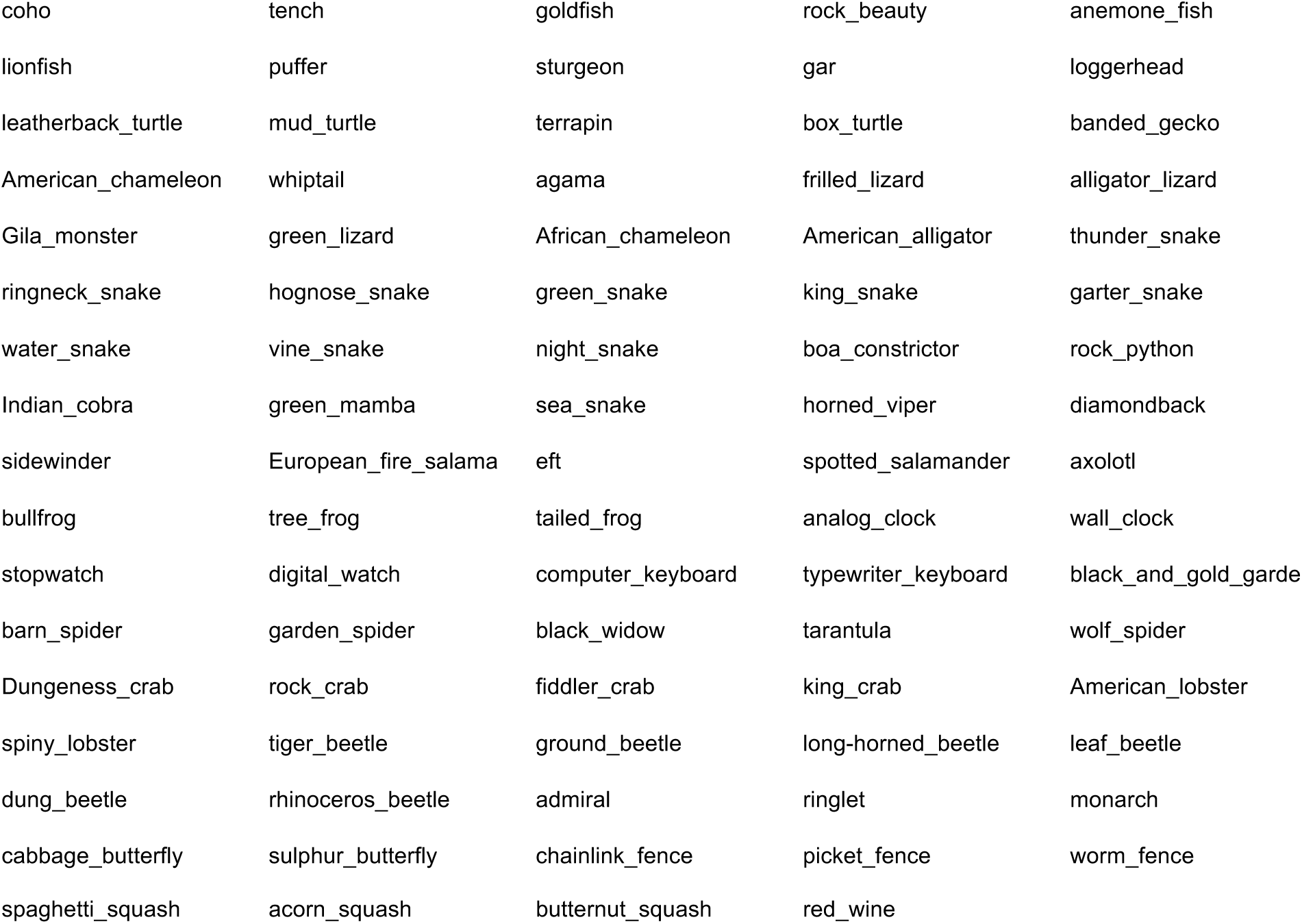
ImageNet categories were manually labeled as either basic- or subordinate-level (see *Methods*). To find the folder ID-to-category name correspondence, see the file “bOrS.csv” in our Github repository (https://github.com/ojinsi/startingblurry).

**Supplementary Table 2.**
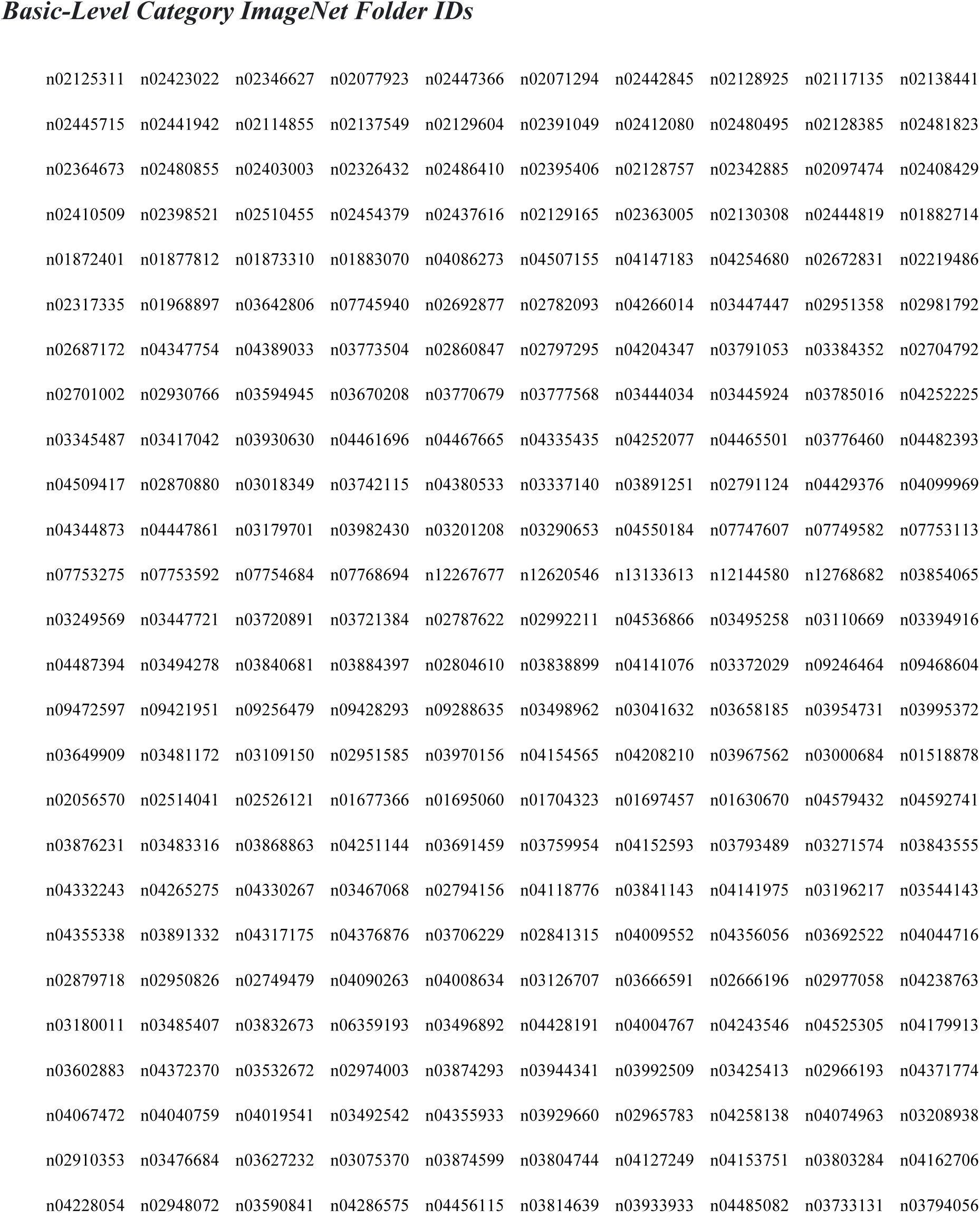
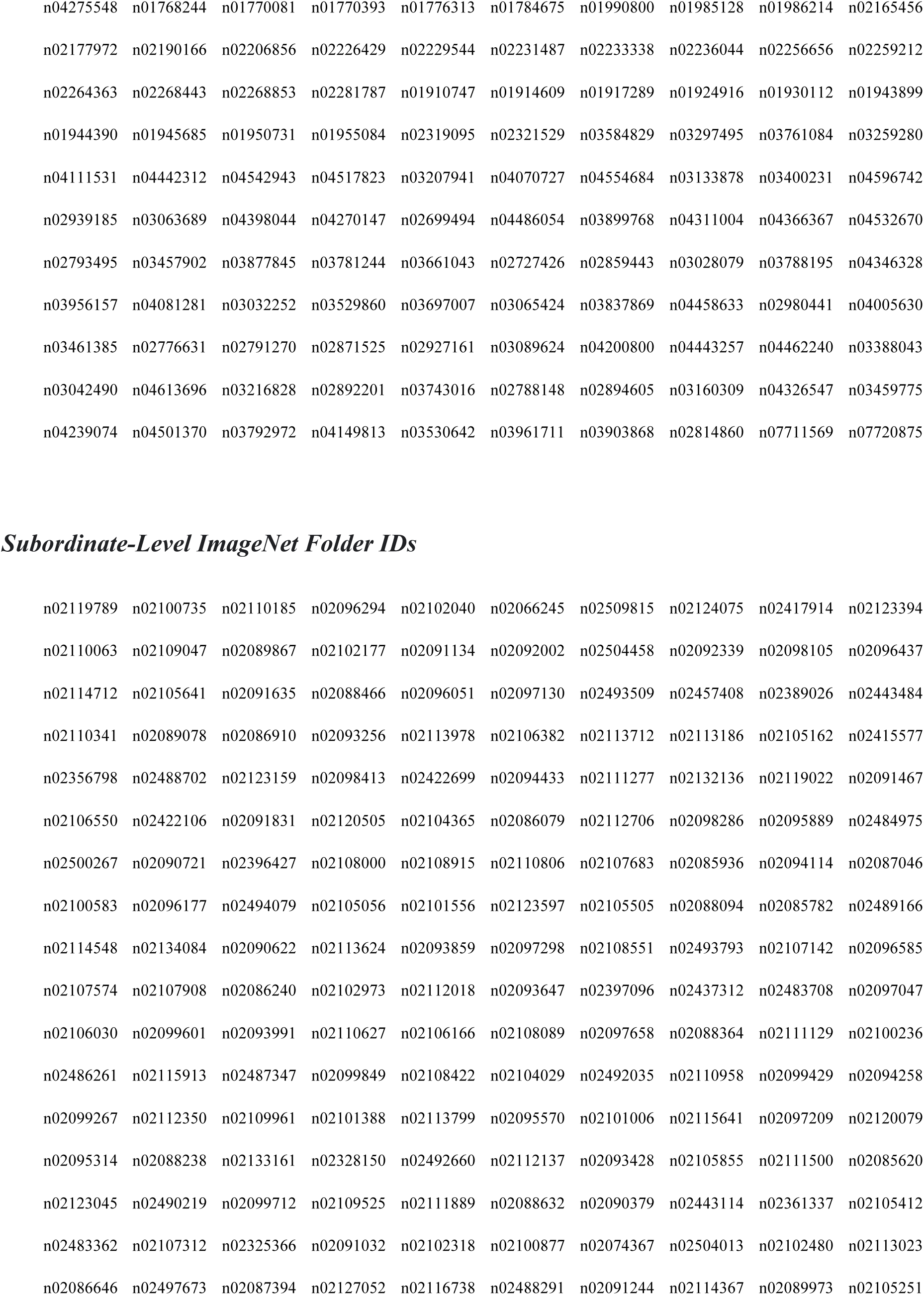
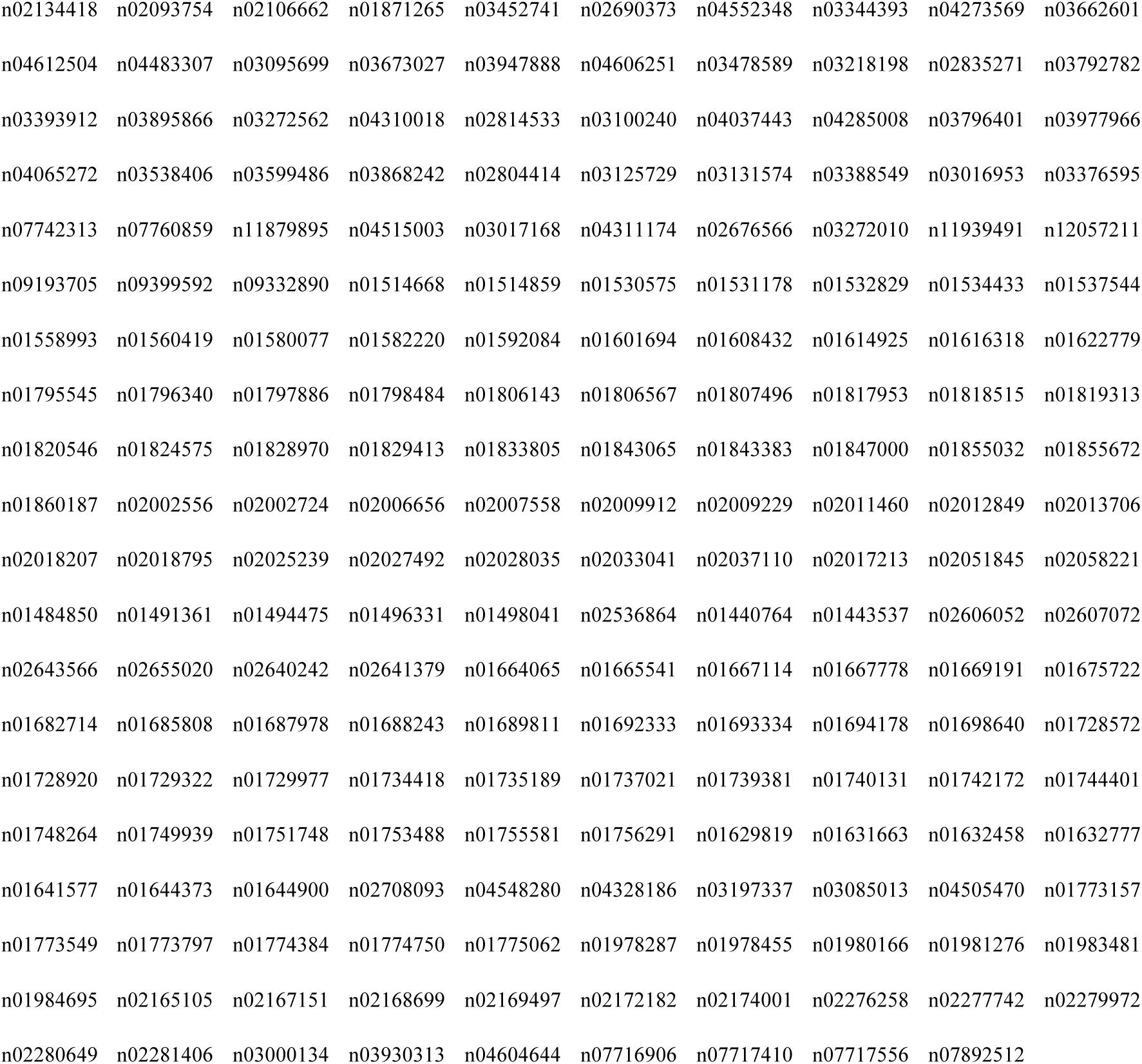
ImageNet categories were manually labeled as either basic- or subordinate-level (see *Methods*). To find the category-to-folder ID name correspondence, see the file “bOrS.csv” in our Github repository (https://github.com/ojinsi/startingblurry).

|  |  |  |  |  |  |  |  |  |  |
| --- | --- | --- | --- | --- | --- | --- | --- | --- | --- |
| n02125311 | n02423022 | n02346627 | n02077923 | n02447366 | n02071294 | n02442845 | n02128925 | n02117135 | n02138441 |
| n02445715 | n02441942 | n02114855 | n02137549 | n02129604 | n02391049 | n02412080 | n02480495 | n02128385 | n02481823 |
| n02364673 | n02480855 | n02403003 | n02326432 | n02486410 | n02395406 | n02128757 | n02342885 | n02097474 | n02408429 |
| n02410509 | n02398521 | n02510455 | n02454379 | n02437616 | n02129165 | n02363005 | n02130308 | n02444819 | n01882714 |
| n01872401 | n01877812 | n01873310 | n01883070 | n04086273 | n04507155 | n04147183 | n04254680 | n02672831 | n02219486 |
| n02317335 | n01968897 | n03642806 | n07745940 | n02692877 | n02782093 | n04266014 | n03447447 | n02951358 | n02981792 |
| n02687172 | n04347754 | n04389033 | n03773504 | n02860847 | n02797295 | n04204347 | n03791053 | n03384352 | n02704792 |
| n02701002 | n02930766 | n03594945 | n03670208 | n03770679 | n03777568 | n03444034 | n03445924 | n03785016 | n04252225 |
| n03345487 | n03417042 | n03930630 | n04461696 | n04467665 | n04335435 | n04252077 | n04465501 | n03776460 | n04482393 |
| n04509417 | n02870880 | n03018349 | n03742115 | n04380533 | n03337140 | n03891251 | n02791124 | n04429376 | n04099969 |
| n04344873 | n04447861 | n03179701 | n03982430 | n03201208 | n03290653 | n04550184 | n07747607 | n07749582 | n07753113 |
| n07753275 | n07753592 | n07754684 | n07768694 | n12267677 | n12620546 | n13133613 | n12144580 | n12768682 | n03854065 |
| n03249569 | n03447721 | n03720891 | n03721384 | n02787622 | n02992211 | n04536866 | n03495258 | n03110669 | n03394916 |
| n04487394 | n03494278 | n03840681 | n03884397 | n02804610 | n03838899 | n04141076 | n03372029 | n09246464 | n09468604 |
| n09472597 | n09421951 | n09256479 | n09428293 | n09288635 | n03498962 | n03041632 | n03658185 | n03954731 | n03995372 |
| n03649909 | n03481172 | n03109150 | n02951585 | n03970156 | n04154565 | n04208210 | n03967562 | n03000684 | n01518878 |
| n02056570 | n02514041 | n02526121 | n01677366 | n01695060 | n01704323 | n01697457 | n01630670 | n04579432 | n04592741 |
| n03876231 | n03483316 | n03868863 | n04251144 | n03691459 | n03759954 | n04152593 | n03793489 | n03271574 | n03843555 |
| n04332243 | n04265275 | n04330267 | n03467068 | n02794156 | n04118776 | n03841143 | n04141975 | n03196217 | n03544143 |
| n04355338 | n03891332 | n04317175 | n04376876 | n03706229 | n02841315 | n04009552 | n04356056 | n03692522 | n04044716 |
| n02879718 | n02950826 | n02749479 | n04090263 | n04008634 | n03126707 | n03666591 | n02666196 | n02977058 | n04238763 |
| n03180011 | n03485407 | n03832673 | n06359193 | n03496892 | n04428191 | n04004767 | n04243546 | n04525305 | n04179913 |
| n03602883 | n04372370 | n03532672 | n02974003 | n03874293 | n03944341 | n03992509 | n03425413 | n02966193 | n04371774 |
| n04067472 | n04040759 | n04019541 | n03492542 | n04355933 | n03929660 | n02965783 | n04258138 | n04074963 | n03208938 |
| n02910353 | n03476684 | n03627232 | n03075370 | n03874599 | n03804744 | n04127249 | n04153751 | n03803284 | n04162706 |
| n04228054 | n02948072 | n03590841 | n04286575 | n04456115 | n03814639 | n03933933 | n04485082 | n03733131 | n03794056 |
| n04275548 | n01768244 | n01770081 | n01770393 | n01776313 | n01784675 | n01990800 | n01985128 | n01986214 | n02165456 |
| n02177972 | n02190166 | n02206856 | n02226429 | n02229544 | n02231487 | n02233338 | n02236044 | n02256656 | n02259212 |
| n02264363 | n02268443 | n02268853 | n02281787 | n01910747 | n01914609 | n01917289 | n01924916 | n01930112 | n01943899 |
| n01944390 | n01945685 | n01950731 | n01955084 | n02319095 | n02321529 | n03584829 | n03297495 | n03761084 | n03259280 |
| n04111531 | n04442312 | n04542943 | n04517823 | n03207941 | n04070727 | n04554684 | n03133878 | n03400231 | n04596742 |
| n02939185 | n03063689 | n04398044 | n04270147 | n02699494 | n04486054 | n03899768 | n04311004 | n04366367 | n04532670 |
| n02793495 | n03457902 | n03877845 | n03781244 | n03661043 | n02727426 | n02859443 | n03028079 | n03788195 | n04346328 |
| n03956157 | n04081281 | n03032252 | n03529860 | n03697007 | n03065424 | n03837869 | n04458633 | n02980441 | n04005630 |
| n03461385 | n02776631 | n02791270 | n02871525 | n02927161 | n03089624 | n04200800 | n04443257 | n04462240 | n03388043 |
| n03042490 | n04613696 | n03216828 | n02892201 | n03743016 | n02788148 | n02894605 | n03160309 | n04326547 | n03459775 |
| n04239074 | n04501370 | n03792972 | n04149813 | n03530642 | n03961711 | n03903868 | n02814860 | n07711569 | n07720875 |
| *Subordinate-Level ImageNet Folder IDs* |  |  |  |  |  |  |  |  |  |
| n02119789 | n02100735 | n02110185 | n02096294 | n02102040 | n02066245 | n02509815 | n02124075 | n02417914 | n02123394 |
| n02110063 | n02109047 | n02089867 | n02102177 | n02091134 | n02092002 | n02504458 | n02092339 | n02098105 | n02096437 |
| n02114712 | n02105641 | n02091635 | n02088466 | n02096051 | n02097130 | n02493509 | n02457408 | n02389026 | n02443484 |
| n02110341 | n02089078 | n02086910 | n02093256 | n02113978 | n02106382 | n02113712 | n02113186 | n02105162 | n02415577 |
| n02356798 | n02488702 | n02123159 | n02098413 | n02422699 | n02094433 | n02111277 | n02132136 | n02119022 | n02091467 |
| n02106550 | n02422106 | n02091831 | n02120505 | n02104365 | n02086079 | n02112706 | n02098286 | n02095889 | n02484975 |
| n02500267 | n02090721 | n02396427 | n02108000 | n02108915 | n02110806 | n02107683 | n02085936 | n02094114 | n02087046 |
| n02100583 | n02096177 | n02494079 | n02105056 | n02101556 | n02123597 | n02105505 | n02088094 | n02085782 | n02489166 |
| n02114548 | n02134084 | n02090622 | n02113624 | n02093859 | n02097298 | n02108551 | n02493793 | n02107142 | n02096585 |
| n02107574 | n02107908 | n02086240 | n02102973 | n02112018 | n02093647 | n02397096 | n02437312 | n02483708 | n02097047 |
| n02106030 | n02099601 | n02093991 | n02110627 | n02106166 | n02108089 | n02097658 | n02088364 | n02111129 | n02100236 |
| n02486261 | n02115913 | n02487347 | n02099849 | n02108422 | n02104029 | n02492035 | n02110958 | n02099429 | n02094258 |
| n02099267 | n02112350 | n02109961 | n02101388 | n02113799 | n02095570 | n02101006 | n02115641 | n02097209 | n02120079 |
| n02095314 | n02088238 | n02133161 | n02328150 | n02492660 | n02112137 | n02093428 | n02105855 | n02111500 | n02085620 |
| n02123045 | n02490219 | n02099712 | n02109525 | n02111889 | n02088632 | n02090379 | n02443114 | n02361337 | n02105412 |
| n02483362 | n02107312 | n02325366 | n02091032 | n02102318 | n02100877 | n02074367 | n02504013 | n02102480 | n02113023 |
| n02086646 | n02497673 | n02087394 | n02127052 | n02116738 | n02488291 | n02091244 | n02114367 | n02089973 | n02105251 |
| n02134418 | n02093754 | n02106662 | n01871265 | n03452741 | n02690373 | n04552348 | n03344393 | n04273569 | n03662601 |
| n04612504 | n04483307 | n03095699 | n03673027 | n03947888 | n04606251 | n03478589 | n03218198 | n02835271 | n03792782 |
| n03393912 | n03895866 | n03272562 | n04310018 | n02814533 | n03100240 | n04037443 | n04285008 | n03796401 | n03977966 |
| n04065272 | n03538406 | n03599486 | n03868242 | n02804414 | n03125729 | n03131574 | n03388549 | n03016953 | n03376595 |
| n07742313 | n07760859 | n11879895 | n04515003 | n03017168 | n04311174 | n02676566 | n03272010 | n11939491 | n12057211 |
| n09193705 | n09399592 | n09332890 | n01514668 | n01514859 | n01530575 | n01531178 | n01532829 | n01534433 | n01537544 |
| n01558993 | n01560419 | n01580077 | n01582220 | n01592084 | n01601694 | n01608432 | n01614925 | n01616318 | n01622779 |
| n01795545 | n01796340 | n01797886 | n01798484 | n01806143 | n01806567 | n01807496 | n01817953 | n01818515 | n01819313 |
| n01820546 | n01824575 | n01828970 | n01829413 | n01833805 | n01843065 | n01843383 | n01847000 | n01855032 | n01855672 |
| n01860187 | n02002556 | n02002724 | n02006656 | n02007558 | n02009912 | n02009229 | n02011460 | n02012849 | n02013706 |
| n02018207 | n02018795 | n02025239 | n02027492 | n02028035 | n02033041 | n02037110 | n02017213 | n02051845 | n02058221 |
| n01484850 | n01491361 | n01494475 | n01496331 | n01498041 | n02536864 | n01440764 | n01443537 | n02606052 | n02607072 |
| n02643566 | n02655020 | n02640242 | n02641379 | n01664065 | n01665541 | n01667114 | n01667778 | n01669191 | n01675722 |
| n01682714 | n01685808 | n01687978 | n01688243 | n01689811 | n01692333 | n01693334 | n01694178 | n01698640 | n01728572 |
| n01728920 | n01729322 | n01729977 | n01734418 | n01735189 | n01737021 | n01739381 | n01740131 | n01742172 | n01744401 |
| n01748264 | n01749939 | n01751748 | n01753488 | n01755581 | n01756291 | n01629819 | n01631663 | n01632458 | n01632777 |
| n01641577 | n01644373 | n01644900 | n02708093 | n04548280 | n04328186 | n03197337 | n03085013 | n04505470 | n01773157 |
| n01773549 | n01773797 | n01774384 | n01774750 | n01775062 | n01978287 | n01978455 | n01980166 | n01981276 | n01983481 |
| n01984695 | n02165105 | n02167151 | n02168699 | n02169497 | n02172182 | n02174001 | n02276258 | n02277742 | n02279972 |
| n02280649 | n02281406 | n03000134 | n03930313 | n04604644 | n07716906 | n07717410 | n07717556 | n07892512 |  |

